# Time-lagged Flux in the Transition Path Ensemble: Flux Maximization and Relation to Transition Path Theory

**DOI:** 10.1101/2022.02.23.481712

**Authors:** Wenjin Li

## Abstract

Transition path ensemble is of special interest in reaction coordinate identification as it consists of reactive trajectories that start from the reactant state and end in the product one. As a theoretical framework for describing the transition path ensemble, the transition path theory has been introduced more than ten years ago and so far its applications have been only illustrated in several low-dimensional systems. Given the transition path ensemble, expressions for calculating flux, current (a vector field), and principal curve are derived here in the space of collective variables from the transition path theory and they are applicable to time-series obtained from molecular dynamics simulations of high-dimensional systems, i.e., the position coordinates as a function of time in the transition path ensemble. The connection of the transition path theory is made to a density-weighted average flux, a quantity proposed in a previous work to appraise the relevance of a coordinate to the reaction coordinate [W. Li, J. Chem. Phys. 156, 054117 (2022)]. Most importantly, as an extension of the existing quantities, time-lagged quantities such as flux and current are also proposed. The main insights and objects provided by these time-lagged quantities are illustrated in the application to the alanine peptide in vacuum.

## Introduction

One of the fundamental problems in the field of computational biochemistry and biophysics is the study of rare events, such as chemical reactions, protein folding, and protein-ligand interactions. These rare events often-times can be simplified as rare transitions between two dominant stable states A and B separated by high activation barrier.^1^ As for chemical reactions, for instance, the stable states A and B correspond to the reactant state and the product state, respectively. When focusing on the dynamic aspects of the rare events, a transition from one state to the other completes in a very short time (named the transition time) that is usually orders of magnitude less than the mean first-passage time. Thus, these rare transitions or transition paths are of vital importance in both theoretical and computational study on complex systems. ^2,3^ As a statistical theory for describing the ensemble of transition paths, the transition path theory pioneered by E and Vanden-Eijnden^2^ shows that the steady state flux is proportional to the number of transition paths or reactive trajectories from A to B. The probability density of the states in the transition path ensemble (TPE) is related to the equilibrium probability density by the committor or splitting probability, the probability that the system starting from a given state will commit to the state B first before it reaches the state A.^4–6^ The reactive probability current is also proposed to be a useful quantity that characterize how transition paths move from A to B on average, and it is given by the product of the equilibrium probability times the gradient of the committor function in case of an overdamped dynamics with isotropic diffusion. If the diffusion is anisotropic, the position-dependent diffusion matrix *D* must be taken into account.^7,8^ Then the transition tube in which most of the probability current is located and the principal curve consisting of the points at which the current intensity on each isocommittor surface is maximized can be determined to further characterize the reaction mechanism of the rare event. The transition path theory is then generalized by Metzner and coworkers^9^ to discrete systems in the context of continuous-time Markov chains.

One main application of the transition path theory is the derivation of the string method,^10–14^ which aims to obtain a curve inside the transition tube to best approximate the principal curve in either the phase space or in the space of collective variables (CVs), which are assumed to be capable of capturing the essential dynamics of the system. For instance, the committor function can be approximated by a function of these CVs.^13^ In the string method with swarms-of-trajectories ^14^ (also named the dynamic string method^8^), it is shown recently that the time-length of the short unbiased trajectories should be longer enough to satisfy the conditions for Markovity within the space of CVs.^15^ Bartolucci and coworkers^16^ proposed a theoretical scheme from which the key quantities of TPT, such as the probability density of reactive trajectories, reactive current, and principal curve, can be estimated using short non-equilibrium trajectories obtained from biased simulations such as self-consistent path sampling and ratchet-and-pawl molecular dynamics.

The transition path theory has been also directly applied to calculate the reactive current for several low dimensional systems,^2,17^ such as the rugged Mueller potential, a three-hole potential, and a double-well potential with dynamics ranging from overdamped to underdamped diffusion. Metzner and coworkers^9^ applied the transition path theory for Markov processes to a long equilibrium trajectory of a biomolecular system, that is the trialanine in vacuum, obtained by molecular dynamics (MD) simulations. In the application, the long trajectory was discretized and then converted to a reversible Markov jump process in a subspace of two pre-defined torsion angles. Johnson and Hummer^8^ calculated the reactive currents and reactive pathways in several model potentials to explore the effects from anisotropy and position dependence of diffusion tensors. These systems can be either analytically solved or long equilibrium trajectories of them are available from computer simulations. Since MD simulation is an important tool for studying complex biomolecular systems, ^18,19^ it is thus of great interest to extend the transition path theory to be applicable to analyse the generally-available MD trajectories of these systems. However, the application of the transition path theory to complexes systems remains difficult and computationally expensive.

Since the computational cost in sampling the TPE is generally less than sampling a long equilibrium trajectory, the TPE is one of the common ensembles of trajectories that are available for complex biomolecular systems.^20^ Apart from the string method, several other methods such as nudged elastic-band method,^21,22^ forward flux sampling, ^23,24^ milestoning,^25–27^ weighted ensemble sampling, ^28^ and transition path sampling (TPS),^3,29–32^ are also representative approaches among many others in sampling transition paths. Starting from an initial transition path, TPS samples new transition paths by randomly moving in the space of transition paths if the two stable states A and B can be well-defined by one or several pre-defined order parameters,^3,29–32^ which are not required to be good order parameters to distinguish the states in the transition regions, i.e., the region in between the states A and B. Importantly, the transition paths obtained from TPS are unbiased trajectories that are properly weighted as in an equilibrium ensemble. TPS has been employed in a variety of applications to complex biomolecular systems.^33–42^ Thus, it should be of great interest if the transition path theory can be applied to the TPE of complex systems. As will be shown below, the new formulas of reactive current and principal curve that are applicable to the TPE are derived from the transition path theory. For many of these formulas, the derivations are more than trivial.

Another difficulty in the application of the transition path theory to complex systems is the high dimensionality of these systems. It is practically infeasible to calculate and visualize the reactive current and the corresponding principal curve in the phase space. Generally, one seeks to project them onto a space of a few CVs. Although the extension of the TPT to the CV space is rather straightforward, it is not trivial to select a proper set of CVs to which the effective dynamics of the system can be preserved, which is closely connected to a very challenging problem, i.e., the identification of reaction coordinates.^43^ From a theoretical viewpoint, the ‘ideal’ reaction coordinate is the committor, which however provides very limited information about the reaction mechanism and is arduous to be computed. For a system with parabolic barriers and diffusive dynamics, the gradient of the committor at the saddle point is parallel to the only positive eigenvector of the matrix VD, while the reactive current is directed to the only positive eigenvector of the matrix DV, ^44,45^ where V and D are the Hessian matrix of the potential of mean force and the diffusion matrix, respectively. If ***e*** denotes the unit vector for the gradient of the committor, the direction of the reactive current is parallel to the vector D***e*** for any diffusive process where the Smoluchowski equation is applicable.^7,8^ From a practical viewpoint, it will be useful to identify a few CVs or physical variables to represent the committor. Ma and Dinner^46^ utilized the committor to train an artificial neural network and select automatically important physical variables from a large pool of candidates with a genetic algorithm. Best and Hummer^38^ examined the probability of configurations located on the isosurfaces of a tested coordinate to be also on a reactive trajectory and the reaction coordinate is proposed to be the one along which the distribution of such a probability is most sharply peaked. Other approaches to identify the reaction coordinate based on the committor includes likelihood maximization method and its extensions,^47–49^ cross-entropy minimization method,^50^ and transition state ensemble optimisation.^51,52^ Without the information of the committor, machine learning methods were employed to identity reaction coordinates for complex biomolecular systems. ^53,54^ On the other hand, the committor can be approximately estimated with affordable efforts. Li and coworkers utilized a fitting procedure to reduce the cost of evaluating the committor for configurations in the TPE.^55^ Algorithms such as iso-committor surfaces calculation, ^56^ non-equilibrium non-parametric analysis,^57^ and Markov state models,^58,59^ were proposed to estimate the committor of configurations in an ensemble of non-equilibrium short trajectories.

To tackle the challenge of reaction coordinate identification, many others paid particular attentions to the TPE, in which each transition paths roughly follow the direction of the reaction coordinate in its journey from the reactant state to the product one. Following such a line of thought, the TPE is thus believed to possess the information enough for unveiling the reaction mechanism and extracting the reaction coordinates.^31,43,52,60^ As a quantity calculated from the TPE by combining the atomic forces on all the atoms and their positional coordinates, the so-called emergent potential energy along a coordinate was recently demonstrated to be quite promising in appraising the relevance of a coordinate to the reaction coordinate.^61^ Li then showed that the correct reaction coordinate can be also identified with the equipartition terms in the TPE, which were estimated from the full coordinates (both position and momentum) of the reactive trajectories.^62^ Later, a different viewpoint was suggested in the flux maximization approach.^63^ Noting that each reactive trajectory in the TPE is actually one unit of flux from the reactant state to the product one and the largest possible net flux along a coordinate is the number of transition paths, which equals the flux along the true reaction coordinate, the flux maximization approach quantifies the relevance of a coordinate to the reaction coordinate with a density-weighted average flux along it and tries to identify a one-dimensional reaction coordinate along which the average flux maximizes. Its application to the alanine dipeptide in vacuum obtained a one-dimensional reaction coordinate that is almost parallel to the gradient of the committor ***e***, although the direction of reactive current vectors in diffusive dynamics is known to be determined by the vector D***e***.^7,8^

In this work, we established a direct connection between the net flux defined in Ref. [^63^] and the transition path theory. From such a connection, we immediately realized that the flux maximization aims to identify a one-dimensional coordinate, which is expected to be a monotonically increasing function of the committor (not the committor itself) and is thus able to reproduce the direction of the gradient of the committor. In addition, we explored the properties of the density-weighted averaged flux from a theoretical viewpoint and found that this quantity is closely related to a newly proposed quantity, that is the time-lagged flux, which then leads to the derivation of time-lagged reactive current and the corresponding principal curve. The introduction of these time-lagged quantities is inspired by noticing that the diffusion matrix appears in the formula of the flux when an effective propagator with lag time is used in Ref. [^15^]. The usual reactive flux and current is then a special case of the time-lagged analogues when the lag time approaches zero.

In the following, we start by briefly review the transition path theory. Next, the transition path theory is then generalized to the space of CVs and a connection to it is made for the net flux in the TPE proposed in Ref. [^63^]. Then, the properties of the density-weighted average flux is explored and the time-lagged flux, current and principal curve are introduced. Thereafter, we illustrate the application of the proposed theoretical framework to the *C*_7eq_ → *C*_7ax_ isomerization of the alanine dipeptide in vacuum, a well-studied system with its reaction mechanisms being largely known.^46,50,61,62,64,65^ We end the paper with concluding remarks.

## Theory

### Probability density and probability current in the transition path ensemble

Given an infinitely long trajectory ***q***(*t*) with ***q*** being a point in the configuration space ***Ω*** and *t* ≥ 0 (here we largely follow the notation and nomenclature in Ref. [^2^]), there exists many reactive trajectories from A to B that start from A and reach B without visiting A in between when we consider a prototypical system consisting of two stable states A and B. These reactive trajectories are also referred as transition paths. An ensemble of reactive trajectories from A to B (also named as A-B reactive trajectories or A-B transition paths in this work) is thus called the TPE from A to B (TPE_*AB*_).

According to the transition path theory,^2^ the probability density of finding a reactive trajectory at ***q*** or the conditional probability density of finding a configuration at ***q*** in the TPE_*AB*_ is

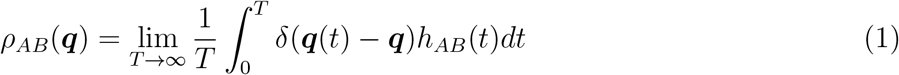

where *h*_*AB*_(*t*) is the indicator function for the TPE_*AB*_, that is *h*_*AB*_(*t*)=1 if ***q***(*t*) belongs to an A-B reactive trajectory and *h*_*AB*_(*t*)=0 otherwise. Note that *ρ*_*AB*_(***q***) is not normalized as the integral of *ρ*_*AB*_(***q***) over the configuration space ***Ω*** gives the fraction of the whole trajectory which consists of A-B transition paths. Analogously, the probability density of finding a configuration at ***q*** is

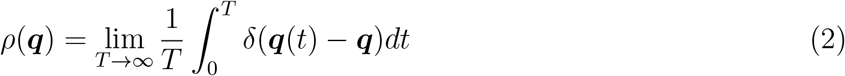

The probability current associated with the A-B transition paths characterizes how these paths move from A to B on average and it is the vector field ***J***_*AB*_ defined as

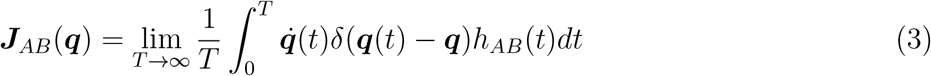

This current is divergence-free in the transition region between A and B, thus the probability flux through any dividing surface between A and B is a constant, that is the reaction rate *v*_*R*_:

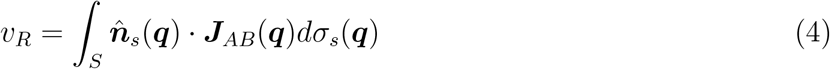

where *dσ*_*s*_(***q***) is the surface element at a point ***q*** in the dividing surface *S*, and 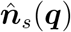 is the unit normal vector to *dσ*_*s*_(***q***) pointing toward B. On the other hand, the reaction rate is related to the number of A-B reactive trajectories (*N*_*T*_) in a given trajectory ***q***(*t*) of finite length *T* by

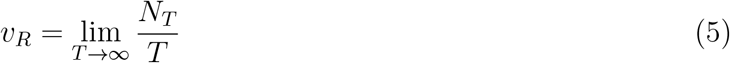

### Flux along a coordinate in the transition path ensemble

From Eqs. (3)-(5), we immediately have

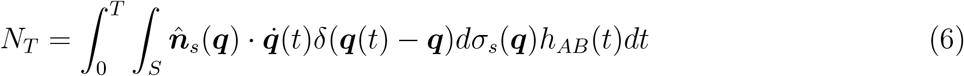

Thus, the net number of reactive or productive transitions across any dividing surface between A and B equals the number of A-B reactive trajectories for a trajectory ***q***(*t*) of arbitrary length. In other words, the net number of transitions across any dividing surface between A and B in the TPE_*AB*_ equals the number of paths in the ensemble. What about the net number of transitions across a dividing surface in the configuration space that is not necessarily a dividing surface between A and B? Let us define by 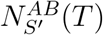 the net number of transitions across a dividing surface *S′* in the configuration space ***Ω*** in the TPE_*AB*_, that is

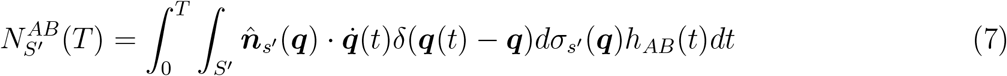

Obviously, 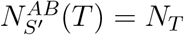 if the dividing surface is also a dividing surface between A and B, otherwise 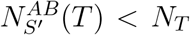. Therefore, the quantity in Eq. 7 can be used to find dividing surfaces between A and B if the TPE_*AB*_ is provided. From now on, we name 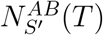 as the flux in the TPE_*AB*_ across a dividing surface. Note that, the flux 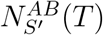 differs from the one obtained with the probability current in Eq. (3) by a normalization factor 1*/T*. When the TPE_*AB*_ is only obtained, the flux 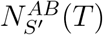 can be estimated although *T* is unknown. Thus, the flux 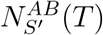 given in Eq. (7) is practically useful in such a case.

We now consider *ξ*(***q***) as a function of ***q*** or a generalized coordinate and *ξ*(***q***) = *ξ′* is a dividing surface in the configuration space, the flux in the TPE_*AB*_ across *ξ*(***q***) = *ξ′* is

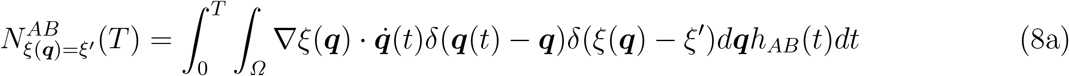

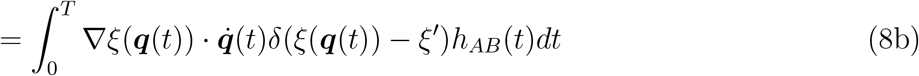

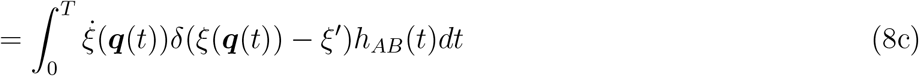

Here we use 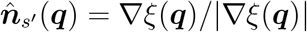 and *dσ*_*s′*_ (***q***) (the surface element in *ξ*(***q***) = *ξ′*) is replaced by *δ*(*ξ*(***q***) − *ξ′*)|∇*ξ*(***q***)|*d****q*** when shifting from surface integral to configuration space integral. Note that the term in the right hand of Eq. 8c is exactly the flux at a particular location *ξ′* of the coordinate *ξ*(***q***) in the TPE_*AB*_ proposed in a previous work.^63^ Thus, we establish the connection for the flux along a coordinate to the transition path theory.

If *ξ*(***q***) is the committor function, one can imagine that 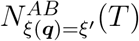 at any value *ξ′* ∈ [0,1] equals *N*_*T*_. It is thus natural to assume that the flux along the reaction coordinate (the committor function is just one solution for the reaction coordinate) equals the number of transition paths in the TPE_*AB*_, the assumption that is used in the previous work^63^ to find a one-dimensional representation of the reaction coordinate. If *ξ*(***q***) is just an approximation of the reaction coordinate, there exists at least one location at which the flux across the surface at this location is less than *N*_*T*_ and there exists certain region in which the flux equals *N*_*T*_, which is also named the plateau region as observed in the flux along the dihedral *ϕ* in the alanine dipeptide in vacuum. If *ξ*(***q***) represents a bath mode, 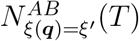 will be vanishingly small everywhere along it, as observed for most of the internal coordinates in the alanine dipeptide in vacuum.^63^ Thus, an variational approach can be designed to find the reaction coordinate. For example, a good reaction coordinate can be identified by finding a coordinate along which the flux is the closest to *N*_*T*_ at any location of it.

So far, we restricted our derivation under the context of the transition path ensemble from A to B. Actually, the integral in Eq. 8c can be performed over an arbitrary set of pieces cut from the long trajectory which is not necessarily the TPE_*AB*_. Let us define by *U* a set of times in [0,T] and *h*_*U*_ (*t*) is the corresponding indicator function, that is *h*_*U*_ (*t*) = 1 if *t* ∈ *U* and *h*_*U*_ (*t*) = 0 otherwise. Thus, *U* is an arbitrary ensemble and the TPE_*AB*_ is a particular example of *U*. Now, we formally define the flux at a give location *ξ′* along a generalized coordinate *ξ*(***q***) in an arbitrary ensemble *U* as

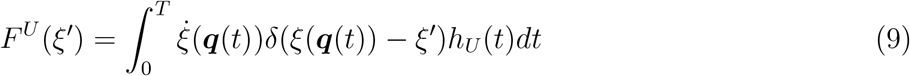

Here, we follow partially the notation in Ref. [^63^] and put *ξ′* inside the bracket to emphasize explicitly that the flux can be considered as a function of *ξ′*. We drop *T* from the bracket of *F*^*U*^ (*ξ′*) by assuming the ensemble U is sufficiently sampled to be a good representative of the one when *T* is infinitely long. The ensemble *U* can be either from a very long straight-forward simulation or it is sampled by enhanced sampling techniques. For example, when *U* stands for the TPE, it can be sampled by the TPS approach.^3,29–32^

Then the density-weighted average flux of a coordinate as defined in Eq. 2 in Ref. [^63^] is

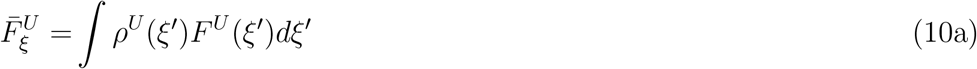

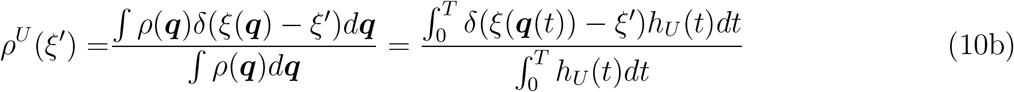

where, *ρ*^*U*^ (*ξ′*) is the normalized probability density of finding a configuration at *ξ′* in the ensemble *U*. Since the 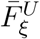 is the weighted average of the flux *F*^*U*^ (*ξ′*) by the probability density, the maximization of 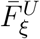 will optimize the region with highest probability density. In the TPE_*AB*_ in the alanine dipeptide in vacuum, the transition state region is mostly populated (Fig. S1), thus the one-dimensional coordinate obtained by maximizing the density-weighted average flux is close to the true reaction coordinate, which is perpendicular to the stochastic separatrix.

Now we consider a different ensemble which consists of the pieces in which the trajectory is going out of A and then returning to A without visiting B, and is thus named as A-A trajectories. These pieces from the trajectory are the trial-and-failed transitions from A to B. We denote the ensemble of such pieces as A-A path ensemble. Similarly, The B-B path ensemble is defined as the collection of the pieces in which the trajectory is going out of B and then returning to B without visiting A. Obviously, the flux across any dividing surface between A and B will vanish in the A-A path ensemble or the B-B path ensemble. We also define the pieces in which the trajectory enters A and after staying in A for a while it leaves A as InA trajectories, and the InB trajectories can be defined analogously.

As shown in Fig. 1, the typical transition path obtained by TPS usually consists of A-B, A-A, B-B, InA, and InB trajectories. Such a path ensemble is named as extended TPE_*AB*_. For the reaction coordinate, e.g., the committor function, the flux along it in the extended TPE_*AB*_ will be the same as the one in the TPE_*AB*_ in the region between A and B. However, the density-weighted average flux obtained from the two ensembles should be different as the probability densities are different due to the inclusion of A-A, B-B, InA and InB trajectories in the extended TPE_*AB*_. Thus, the above analysis suggests that coordinates closely related to and a reasonable one-dimensional representation of the reaction coordinate can be identified by calculating the flux *F*^*U*^ (*ξ′*) and the density-weighted average flux 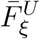 in the extended TPE_*AB*_, as demonstrated in the previous work.^63^

**Figure 1:**
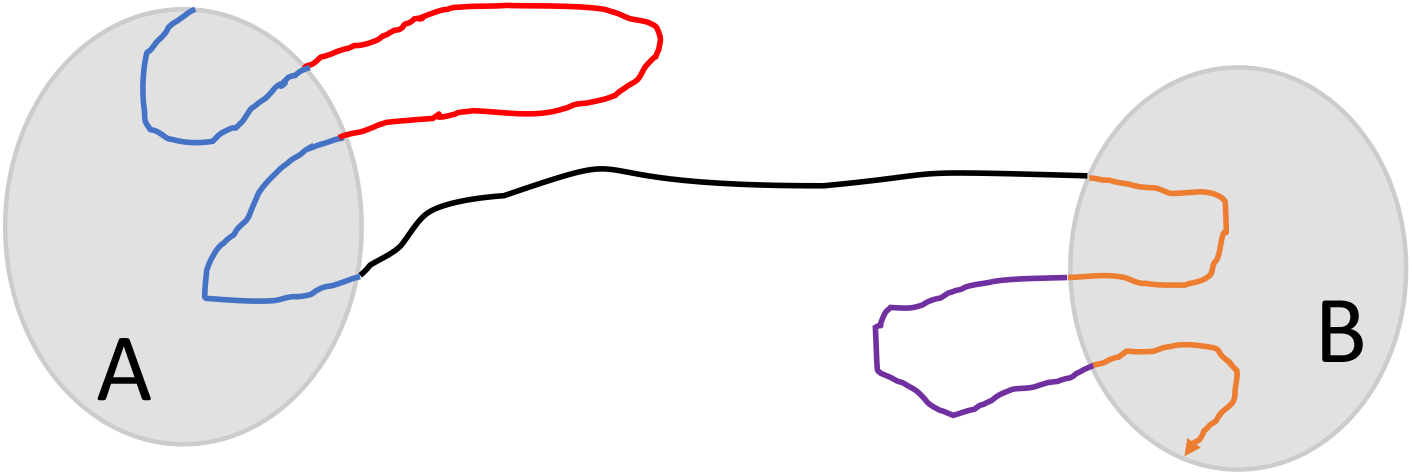
(Schematic) A hypothetical transition path and its various pieces. The reactive piece or an A-B transition path, during which the system starts from the reactant state A and directly reach the product state B, is shown in black. The A-A, B-B, InA, and InB trajectories are shown in red, purple, cyan, and brown, respectively. The definitions of these pieces can be found in the main-text.

One may realize that *F*^*U*^ (*ξ′*) = *F*^*U*^ (*f* (*ξ′*)) and 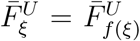 if *f* (*ξ′*) is a monotonically increasing function of *ξ′* (see Appendix A). These properties are practically useful. By maximizing the density-weighted average flux, we aim to find a one-dimensional coordinate of which the committor is a monotonically increasing function. This should be less challenging than directly identifying the committor function itself.

Following the definition of flux in Eq. (9), a quantity that resembles the transmission coefficient can be defined as

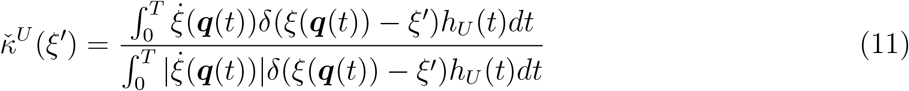

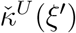 is the ratio of the net number of transitions to the total number of transitions across the dividing surface *ξ*(***q***) = *ξ′* in the ensemble *U*, while the transmission coefficient *κ* is the ratio of the net number of transitions to all the forward crossing events of the surface. Thus, 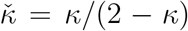 holds.

### Current and principal curves in the collective variable space

For complex systems, it could be impossible to find a reasonable one-dimensional representation of the reaction coordinate. Instead, a set of CVs is usually employed to represent the effective dynamics of the system. These CVs could be identified with the above-mentioned flux calculations, for instance, by selecting a set of top-ranked coordinates according to their density-weighted average flux in the TPE_*AB*_. Let us denote ***y*** ≡ *{y*_1_, *y*_2_, · · ·, *y*_*n*_*}* the set of CVs with *n* being much smaller than the degree of freedom of the system and *y*_*i*_ being a function of ***q***. In analogous to the probability current in Eq. 3 in the TPE_*AB*_, the current in the space of CVs in an arbitrary ensemble *U* is given by

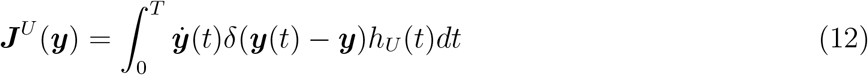

It is a vector field in the space of CVs. Following similar derivations to the ones in the Appendix A, it can be demonstrated that ***J***^*U*^ (***y***) keeps invariant if *y*_*i*_ is replaced by a monotonically increasing function of it. The current in the CVs space is the projection onto it of a non-normalized current in the configuration space (see Appendix B). In case that the number of CVs is two, the vector field can be visualized and one can easily figure out how the reactive paths flow from A to B on average. As suggested by E and Vanden-Eijnden,^2^ the flow lines of this current can be obtained by solving the following artificial dynamics:

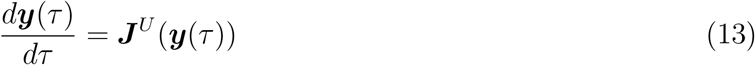

Again, the current ***J***^*U*^ (***y***) differs from the probability current in the same space of CVs by a factor of 1*/T*. It is obvious that the flow lines constructed with Eq. (13) are invariant if ***J***^*U*^ (***y***(*τ*)) is multiplied by a constant. Thus, these flow lines should be the same as the ones constructed from the probability current that is for example obtained from an infinitely long trajectory. These flow lines, which are *n*-dimensional curves in the space of CVs, can be used as one-dimensional representations of the reaction coordinate if they can be properly parametrized such that a dividing surface can also be properly defined at a fixed point along a flow line. The flow lines that carry a dominant portion of the flux form the transition tube that contains most of the reactive current from A to B. It could be useful to identify the flow line which carry high current intensity. The current intensity assigned to a flow line can be the current intensity at its intersection with a given dividing surface, e.g., the isocommittor surface with the committor of 0.5 or simply the dividing surface defined by a generalized coordinate at the free energy barrier top. Alternatively, the average current intensity carried by a flow line can be used and it is given by

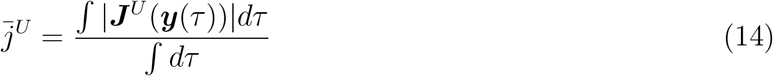

Furthermore, under the localized transition tube assumption, E and Vanden-Eijnden^2^ suggested a procedure to construct the principal curve from the current, which is believed to be a reasonable one-dimensional representation of the reaction pathway. A similar principal curve can be constructed if a generalized coordinate *s*(***y***) is provided, which could be for example the coordinate by maximizing the density-weighted average flux defined in Eq. 10. The principal curve consists of points in the isosurfaces of *s*(***y***) which are centroidal with respect to the current intensity and its point ***η***(*s*) on the surface *s*(***y***) = *s* can be obtained by

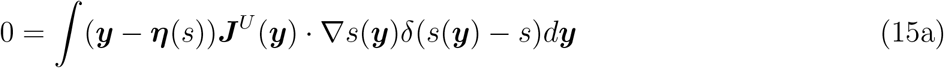

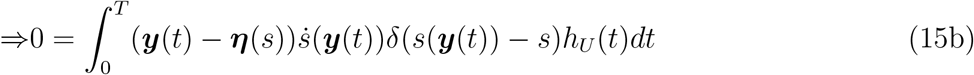

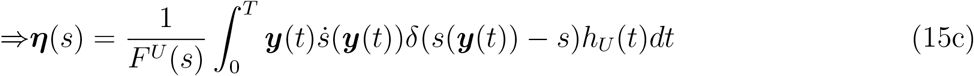

### Time-lagged current and principal curves

As will be seen in the illustrative calculations in the alanine dipeptide in vacuum, the directions of current vectors (Eq. 12) at the transition state region or the barrier top in the plane of *ϕ* and *θ* are different to the reaction coordinate obtained by maximizing the density-weighted average flux (Eq. 10), which on the other hand was shown to be almost perpendicular to the stochastic separatrix.^63^ It is thus of interest to explain such phenomenon from a theoretical viewpoint. Substituting *F*^*U*^ (*ξ′*) in Eq. (9) into Eq. (10) we obtain

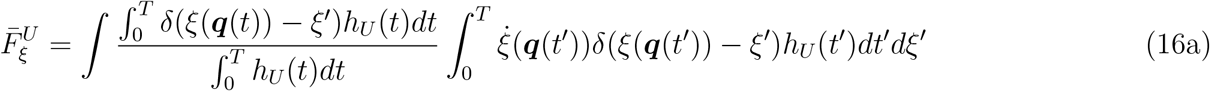

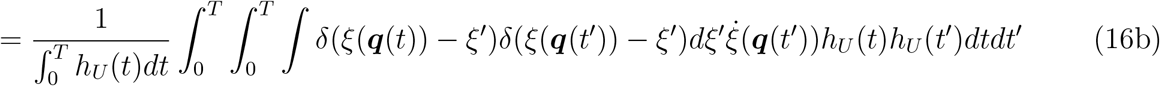

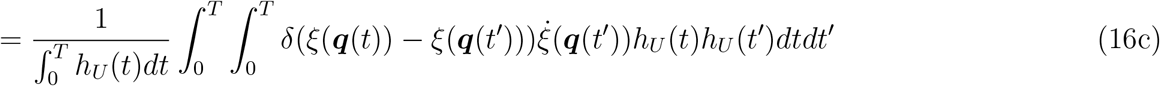

Note that *T* is infinity long and replacing *t′* with *t* + *τ′*, 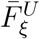 can be further rewritten as

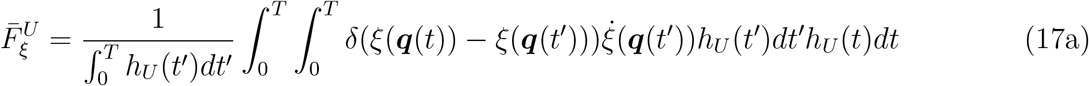

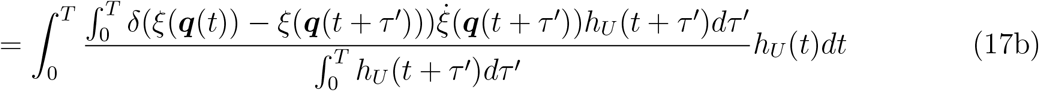

Actually, 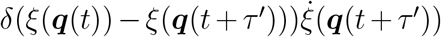 is a time correlation function and 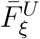 can be viewed as a summation over the averages of this time correlation function. Now, we define a function 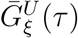 as

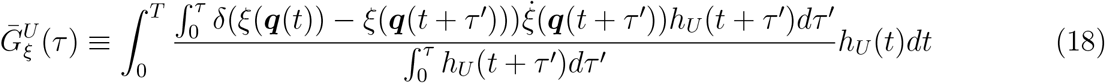

One immediately realize

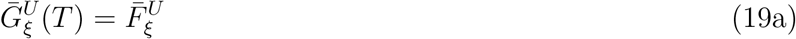

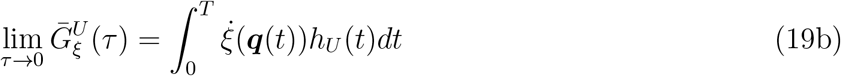

When *U* is chosen as the TPE_*AB*_, then 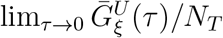 gives the average distance between the boundaries of states A and B along *ξ*. We further define *G*^*U*^ (*ξ′, τ*) as

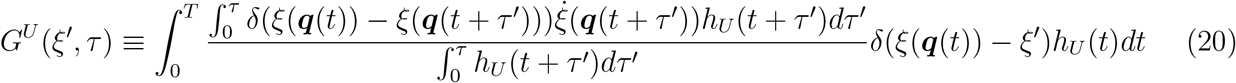

Similarly, we have

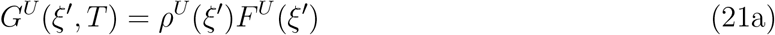

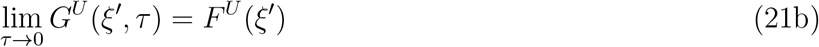

Assuming that the diffusion along *ξ* is slow such that *ξ*(***q***(*t* + *τ′*) is close to *ξ*(***q***(*t*)) within a short time of *τ*, we write 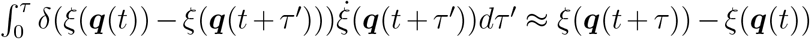. We also assume that the system remains in the ensemble *U* in this short time of *τ* given that it is in *U* at time *t*, thus *h*_*U*_ (*t* + *τ′*) ≈ *h*_*U*_ (*t*). Following through with the above two assumptions yields

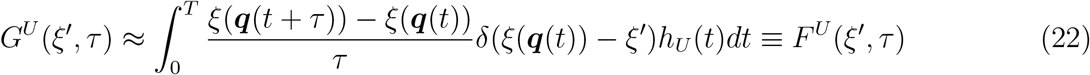

Here the time-lagged flux *F*^*U*^ (*ξ′, τ*) along *ξ* is thus defined. Similarly, we define the time-lagged current ***J***^*U*^ (***y***, *τ*) and the principal curve related to *ξ* in the space of CVs as follows

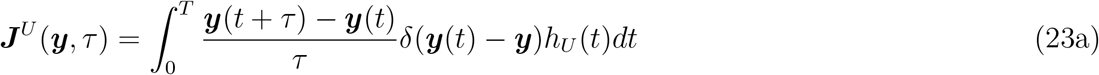

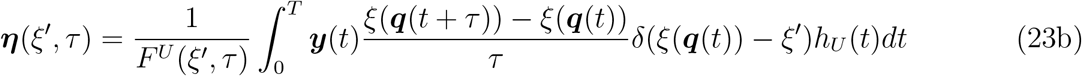

### Differences to the flux-flux correlation function

The terms such as time correlation function and flux above-mentioned can easily remind us a well-known quantity, that is the flux-flux correlation function, ^66–69^ which is mainly used in the computation of reaction rates. We would like to emphasize that the time-lagged flux *F*^*U*^ (*ξ′, τ*), *G*^*U*^ (*ξ′, τ*), and 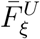 are not reminiscent of the classical flux-flux correlation function. The classical expression of the flux-flux correlation function appeared in the existing literatures is an integral over the phase space with respect to a dividing surface or two different dividing surfaces, it can be expressed as a time integral as below,

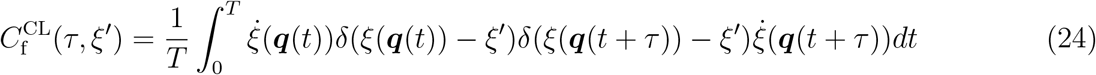

here, the flux-flux correlation function is defined for the dividing surface at *ξ*(***q***) = *ξ′*. Obviously, the classical expression for the flux-flux correlation function in Eq. (24) is different to the ones for the time-lagged flux *F*^*U*^ (*ξ′, τ*) (Eq. (22)), *G*^*U*^ (*ξ′, τ*) (Eq. (20)), and the density-weighted average flux 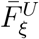 (Eq. (16)). The main differences are listed explicitly as follows: (1) There involves two generalized velocity at time *t* and *t* + *τ* in Eq. (24), while there is just one in the others; (2) The classical flux-flux correlation function is defined for a long equilibrium trajectories, while the others for a non-equilibrium ensemble of trajectory pieces.

### Properties at characteristic lag times

Can we find a value of *τ* at which *F*^*U*^ (*ξ′, τ*) = *G*^*U*^ (*ξ′, T*) or *F*^*U*^ (*ξ′, τ*) resembles *G*^*U*^ (*ξ, T*)? Apparently, lim_*τ*→0_ *F*^*U*^ (*ξ′, τ*) = *F*^*U*^ (*ξ′*). When *τ* approaches infinity, we arrive

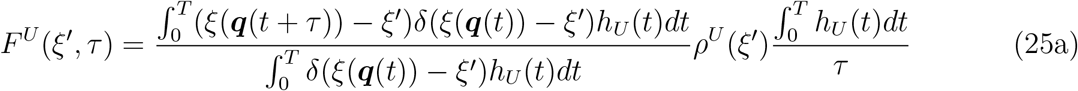

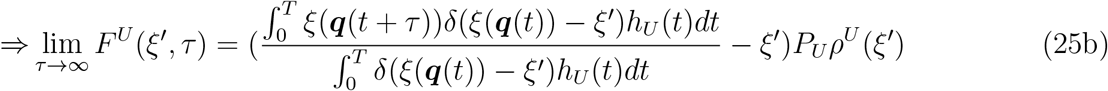

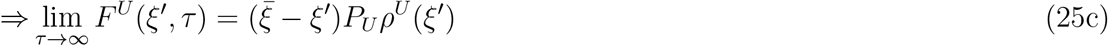

Here, *P*_*U*_ is the fraction of the ensemble *U* in the whole trajectory and 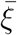 is the average of *ξ*(***q***(*t*)) over the whole trajectory as the distribution of *ξ*(***q***(*t* + *τ*)) will approaches the equilibrium one (when *τ* approaches infinity) given that it was in a fixed location along *ξ* at the time *t*. Although lim_*τ*→0_ *F*^*U*^ (*ξ′, τ*) is much smaller than *G*^*U*^ (*ξ′, T*), both of them have a simple relationship with *ρ*^*U*^ (*ξ′*), thus, it is better to compare the similarity between the functions *F*^*U*^ (*ξ′, τ*) and *G*^*U*^ (*ξ′, T*). We thus consider a particular function 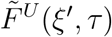

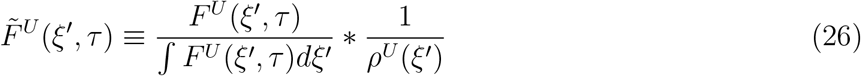

To simply the notations, we define

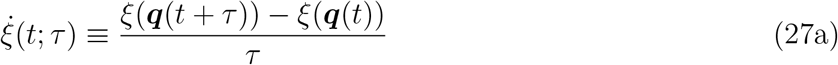

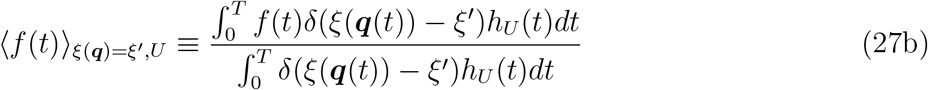

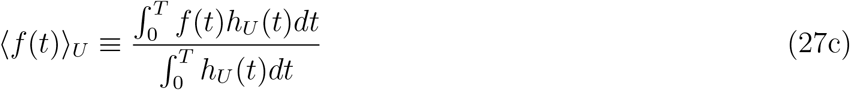

where, *f* (*t*) is an arbitrary function, ⟨· · ·⟩_*ξ*(***q***)=*ξ′,U*_ stands for the conditional average over a surface *ξ*(***q***) = *ξ′* in the ensemble *U*, and ⟨· · ·⟩_*U*_ stands for the average over the ensemble *U*. From Eq. (25c) and (26), after several transformations we have

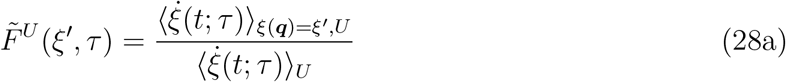

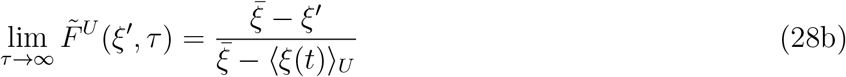

Actually, 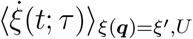 can be viewed as the average velocity of *ξ* at *ξ*(***q***) = *ξ′* in the ensemble *U* when the lag time of an effective dynamical propagator is *τ*. 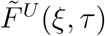 is then the ratio of the average velocity at a particular location along *ξ* to the overall average velocity of the ensemble *U* (shorted as the ratio of average velocity). For a typical system with two stable states separated by a high energy barrier, the transition time from one state to the other (*τ*_*TP*_) is generally far less than the relaxation time *τ*_*rxn*_. Starting from a configuration ***q***(*t*) in a reactive trajectory, the system will commit to the state B after an intermediate time *τ* (i.e., *τ*_*TP*_ *< τ* ≪ *τ*_*rxn*_) by following the reactive trajectory. In another word, ***q***(*t* + *τ*) belongs to the state B if ***q***(*t*) is a point in the extended TPE_*AB*_ for *τ*_*TP*_ *< τ* ≪ *τ*_*rxn*_. If *ξ*(***q***) is a good one-dimensional reaction coordinate and 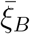 is the average value of *ξ*(***q***) in the state B, we have

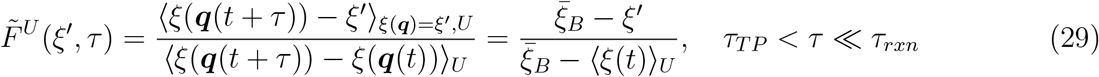

Thus, 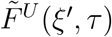 in the intermediate time region is a linear function of *ξ′* with two characteristic points 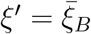 and *ξ′* = ⟨*ξ*(*t*) ⟩_*U*_, at which 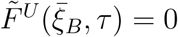 and 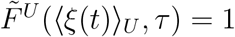, respectively.

Since *G*^*U*^ (*ξ′, T*)*/ρ*^*U*^ (*ξ′*) = *F*^*U*^ (*ξ′*) (Eq. (21a)) and *F*^*U*^ (*ξ′*) is expected to be a constant if *ξ*(***q***) is the true reaction coordinate (e.g., the committor), we thus ask whether it is possible that 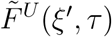 is a constant for a particular *τ* ∗. If *τ* ∗ exists, then the following necessary condition holds

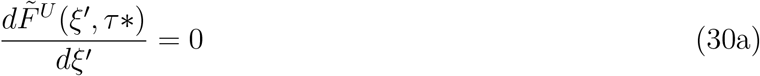

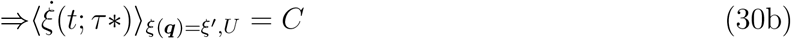

where, *C* is a constant. It is easy to show that 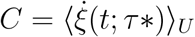 and 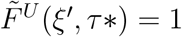. The independence of the average velocity to the location along *ξ* could be an indication of a homogeneous diffusion along *ξ* for a lag time *τ* ∗, thus the time-lagged current vectors in the space of CVs for the lag time *τ* ∗ are believe to be parallel to the gradient of the isocommittor surface. This actually provides a plausible explanation for the observation that the reaction coordinate optimized by flux maximization is perpendicular to the stochastic separatrix, as for a lag time *τ* ∗ the time-lagged flux *F*^*U*^ (*ξ′, τ*) resembles *G*^*U*^ (*ξ′ T*), which is closely related to 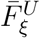 (Eq. (19a)).

One may ask when will *F*^*U*^ (*ξ′, τ*) reach an extreme value, of which the necessary condition is

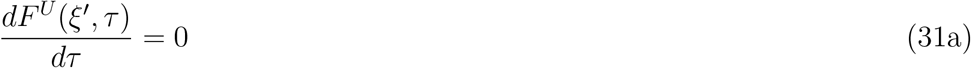

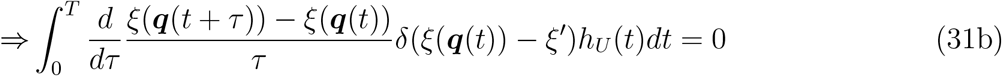

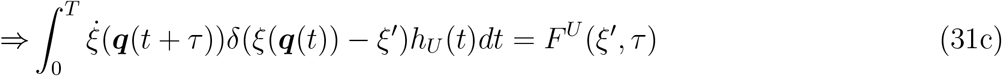

Note that the term on the right hand side of Eq. (31c) is another kind of time-lagged flux and is named 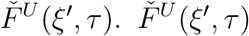 may be regarded as a time-lagged instantaneous flux, while *F*^*U*^ (*ξ′, τ*) is a time-lagged average flux. The time-lagged average flux cross a dividing surface meets its extreme value when it equals the corresponding instantaneous flux.

Similarly, the time-lagged transmission coefficient analogue is defined as

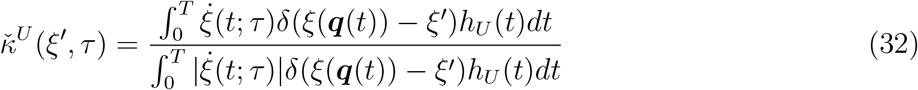

Actually, *τ* is not assumed to be positive. When *τ* is negative, *F*^*U*^ (*ξ′, τ*) could be regarded as a time-lagged backward flux (with an opposite sign) if one names *F*^*U*^ (*ξ′, τ*) as the time-lagged forward flux in case of a positive *τ*. Formally, the time-lagged backward current 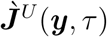 and the time-lagged mean current 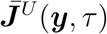 can be defined as

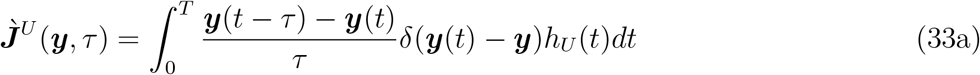

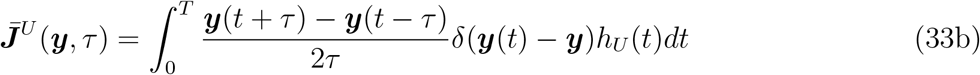

The backward and mean fluxes corresponding to the above currents can also be obtained and their characteristics can be analysed similarly by following the related equations for the time-lagged forward flux *F*^*U*^ (*ξ′, τ*).

Eqs. (22), (23), and (28a) are the central results of the present work and all these quantities can be calculated from the time-series derived from the MD trajectories.

## Illustrative Calculations

The calculation of current and principal curve from a TPE is illustrated with the *C*_7eq_ → *C*_7ax_ isomerization of the alanine dipeptide in vacuum, a well-studied process whose reaction coordinates are known to be two dihedrals *ϕ* and *θ*.^46,50,61–65^ It was shown that the gradient of the committor ***e*** at the transition state region is parallel to a vector 0.78***ϕ*** + 0.63***θ*** (*α* = 39^*°*^).^61^ Here, ***ϕ*** and ***θ*** are the unit vectors along *ϕ* and *θ*, respectively. We use *α* to denote the angle of a vector in the plane of *ϕ* and *θ* to the unit vector ***ϕ*** and *α* ∈ [−180^*°*^, 180^*°*^]. We used a TPE with 5786 transition paths obtained under the same simulation setup as the one in a previous work^63^ and the same definitions are adopted for the sub-ensembles of transition paths, in which the boundaries of state A and B is either defined based on the committor or based on two backbone dihedrals (*ϕ* and *ψ*) and the corresponding sub-ensembles are named with the prefix of “PB” or “PHI”, respectively (See Ref. [^63^] for more details on dividing a TPE into sub-ensembles and their definitions). We first calculated with Eq. (12) the current projected onto the 2D plane of *ϕ* and *θ* for the sub-ensemble *PB*0, which consists of only A-B reactive trajectories with the boundaries of A and B being defined with the committor, and several flow lines. As shown in Fig. 2, almost all the current vectors are pointing in the same direction and the magnitudes of current vectors in the middle are the largest ones, similar properties are also observed for the five flow lines (the probability density of A-B reactive trajectories in the sub-ensemble *PB*0 is shown in Fig. S1). Thus, all the transition paths are located in a localized transition tube. As an estimation of the direction of the current vectors and flow lines, we averaged the directions of the current vectors in a region with *ϕ* ∈ [−7, 7] and *θ* ∈ [−4, 4] and found that the average direction is parallel to a vector 0.92***ϕ*** − 0.39***θ*** (*α* = −23^*°*^), which differ by 62^*°*^ to the vector ***e***. It is not surprising to observe that the directions of current vectors are different to the vector ***e***, as they are determined by *D*(***q***)***e***(***q***) if assuming that the dynamics can be described by the Smoluchowski equation.^7,8^

**Figure 2:**
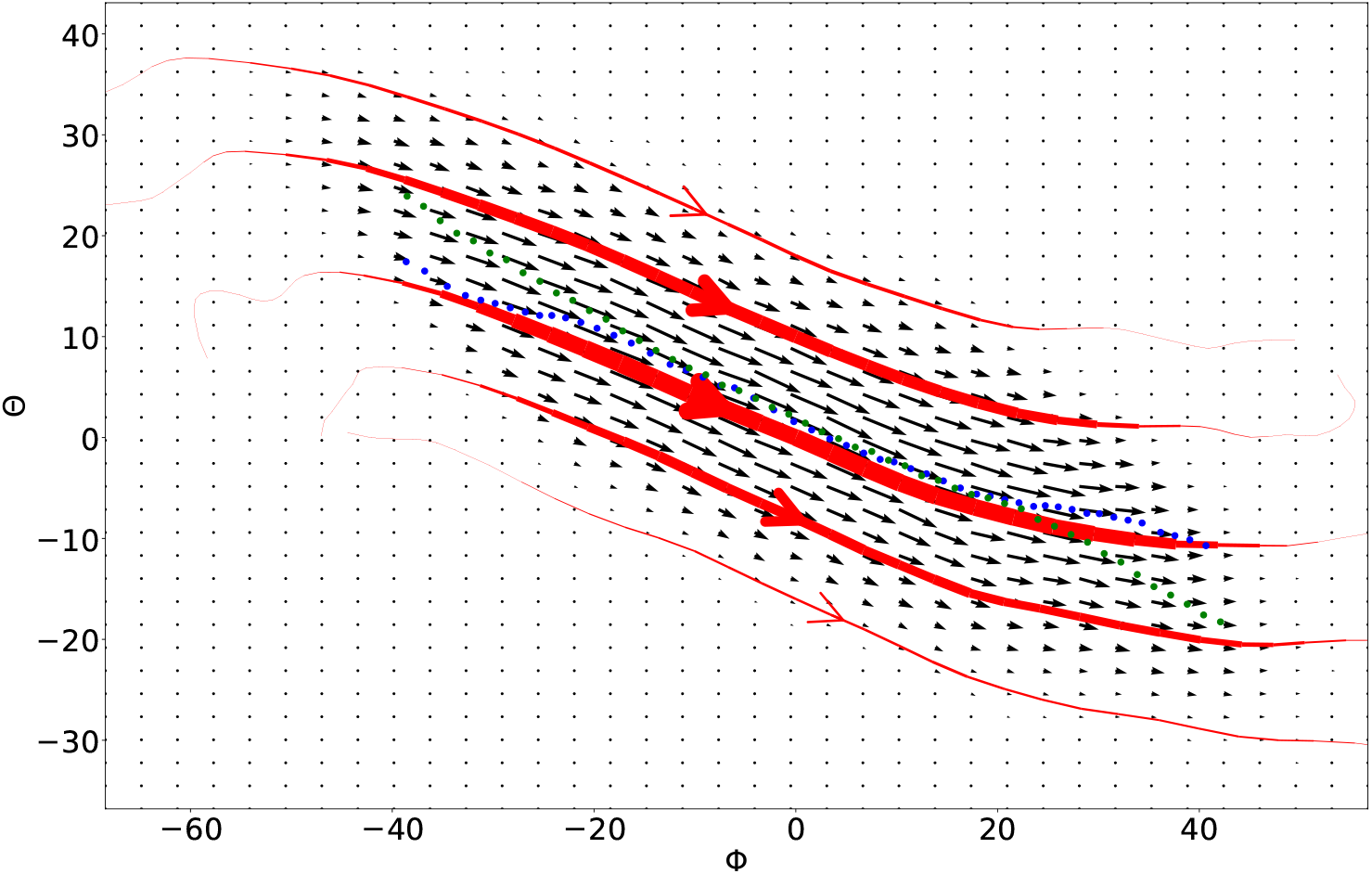
The current, flow lines and principal curves in the plane of two dihedrals *ϕ* and *θ* calculated over the sub-ensemble *PB*0. The current vector field is shown as arrows in black. Five flow lines in red pass the points [0,-16], [0,-8], [0,0], [0,10], and [0,18] on the surface *ϕ* = 0. The width of a flow line varies with the magnitude of the current vector. Blue and green dotted lines are two principal curves obtained with a coordinate *RC*^*opt*^ = 0.83*ϕ* + 0.56*θ* and *ϕ* as the coordinate *s*(***y***), respectively. The units of both axes are degree.

Two principal curves are calculated with Eq. (15c) using a coordinate *RC*^*opt*^ = 0.83*ϕ* + 0.56*θ* and *ϕ* as the coordinate *s*(***y***) (blue and green dotted lines in Fig. 2), respectively. These two principal curves largely coincide, especially in the plateau region (*ϕ* ∈ [−20, 20], Fig. S2) where the flux along *ϕ* equals the number of transition paths, indicating that the location of the principal curve in a region is largely independent on the choice of the generalized coordinate *s*(***y***) if *s*(***y***) defines a good dividing surface in this region. The direction of the principal curve in the region of the barrier top (i.e., *ϕ* ∈ [−15, 10]) is estimated to follow a vector 0.90***ϕ*** − 0.44***θ*** (*α* = −26^*°*^) and it follows closely the direction of the reactive current.

We then projected the current onto two backbone dihedrals *ϕ* and *ψ* (Fig. 3). The current vector and the flow line are pointing along more or less the same direction, as is the case in the *ϕ* and *θ* plane. The width of flow lines are closer compared to the ones in the plane of *ϕ* and *θ*, indicating a broad and uniform distribution of the flux along the transition tube. The two principal curves obtained with the coordinates *RC*^*opt*^ and *ϕ* largely overlap as well and their directions almost coincide to the ones of flow lines, which are parallel to a vector 0.94***ϕ*** − 0.36***ψ***.

**Figure 3:**
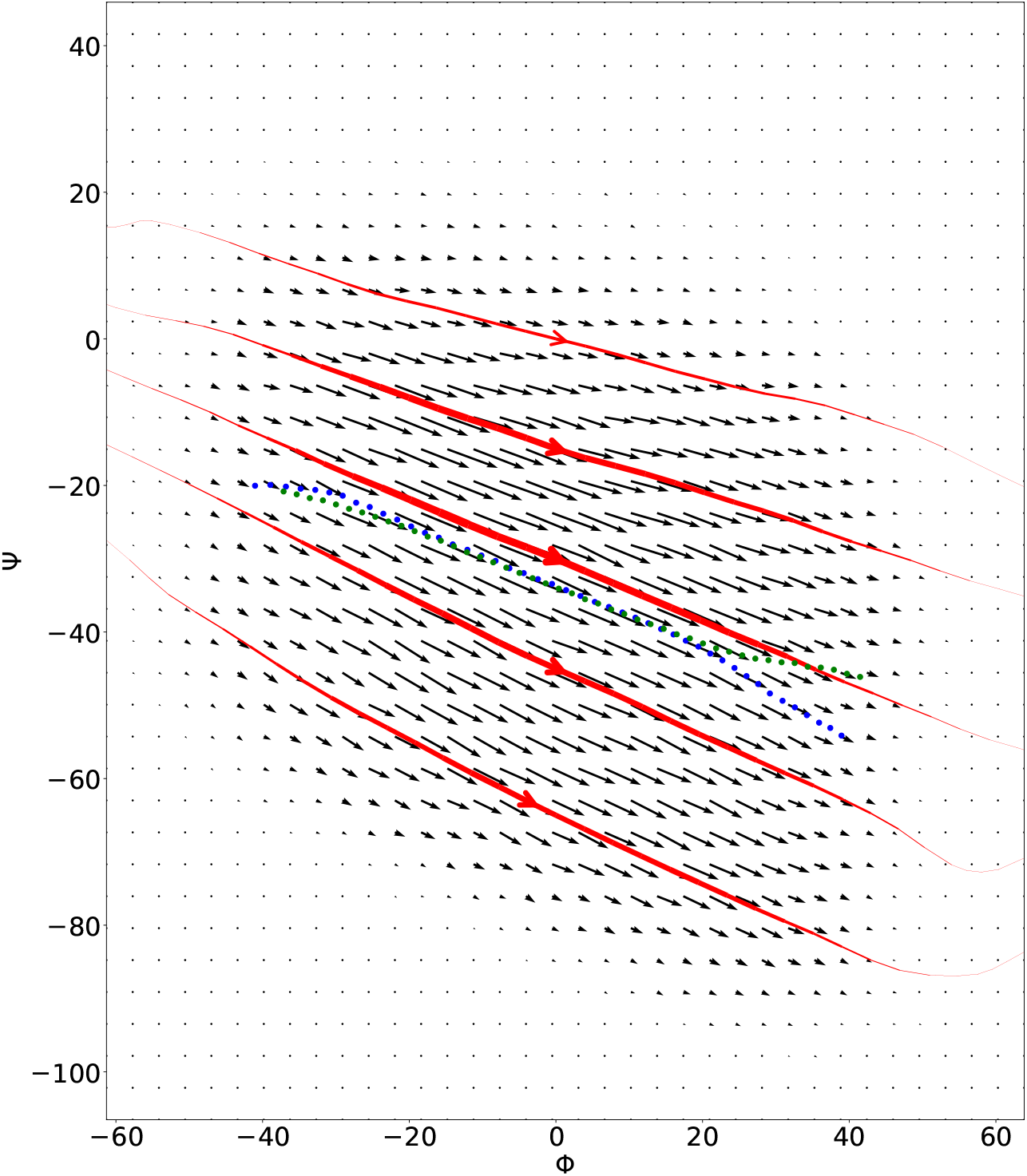
The current, flow lines and principal curves in the plane of two dihedrals *ϕ* and *ψ* calculated over the sub-ensemble *PB*0. The five flow lines pass the points [0,-65], [0,-45], [0,-30], [0,-15], and [0,0] on the surface *ϕ* = 0.

To explore the effects from the definitions of stable basins, similar analyses were performed on an ensemble of reactive trajectories from A to B for which the boundaries of A and B were defined based on *ϕ* and *θ*, that is the sub-ensemble *PHI*0.^63^ As shown in Fig. 4, the flow lines and current vectors are pointing to the same direction to a great extent. The average direction, as obtained by averaging the current vectors in a region with *ϕ* ∈ [−3, 5] and *θ* ∈ [−4, 5], is parallel to a vector 0.92***ϕ*** − 0.39***θ*** (*α* = −23^*°*^), the same as the one obtained with the sub-ensemble *PB*0. To further check the effects from the choice of sub-ensembles, such analyses were carried out on the sub-ensemble *PB*-10 as well, in which the A-A, B-B, InA, and InB trajectories are also included compared to the sub-ensemble *PB*0 (Fig. 5). As observed in Fig. 5, the flow lines remain in the transition tube and the average direction of the reactive current around the barrier top is parallel to a vector 0.93***ϕ*** − 0.37***θ*** (*α* = −22^*°*^), which is almost identical to the ones in the case of sub-ensembles *PB*0 and *PHI*0. In all the sub-ensembles, the directions of the reactive current are almost tangent to the principal curve. Thus, the inclusion of the A-A, B-B, InA, and InB trajectories does not change the direction of reactive current.

**Figure 4:**
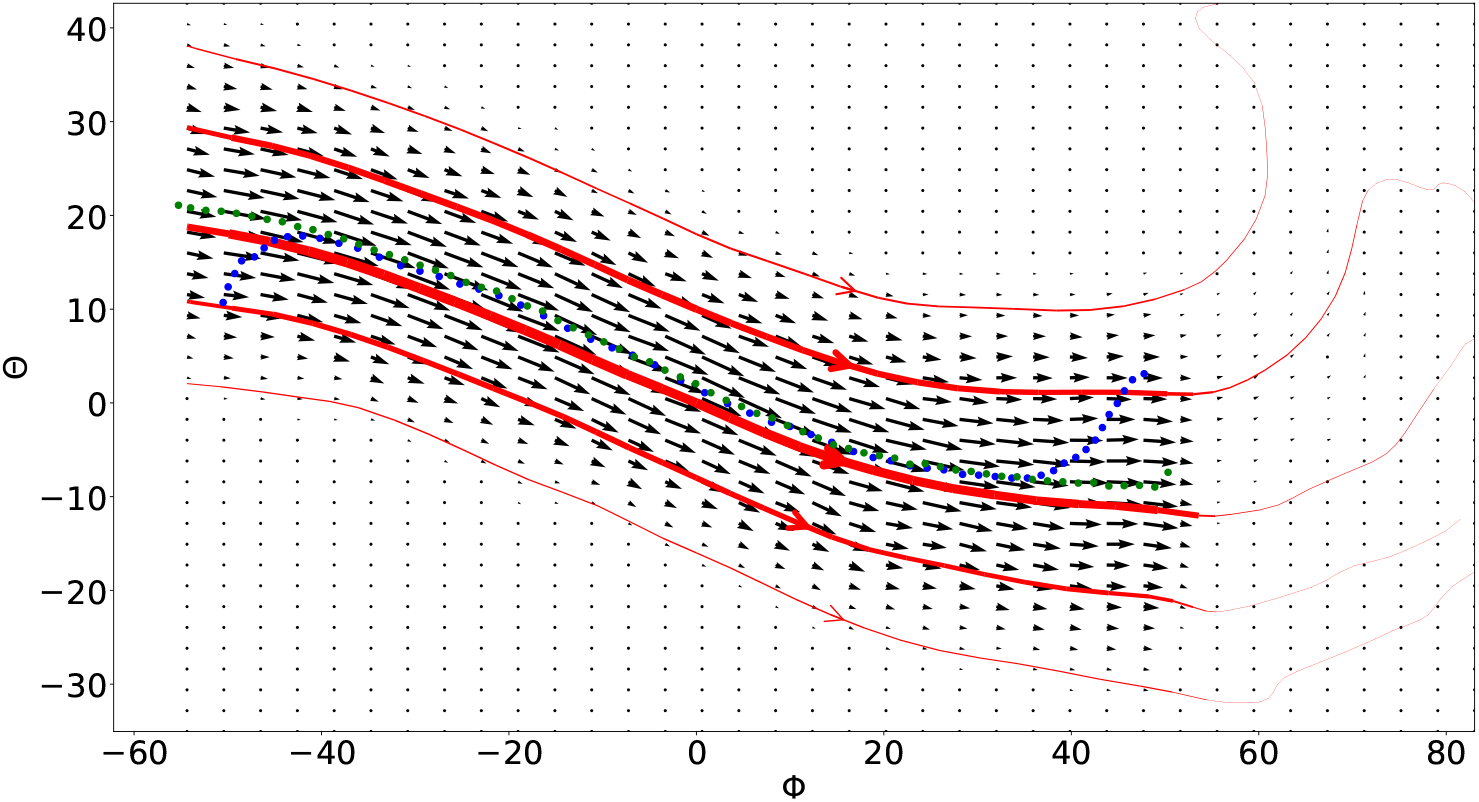
The current, flow lines and principal curves in the plane of two dihedrals *ϕ* and *θ* calculated over the sub-ensemble *PHI*0. The five flow lines pass the points [0,-16], [0,-8], [0,0], [0,10], and [0,18] on the surface *ϕ* = 0.

**Figure 5:**
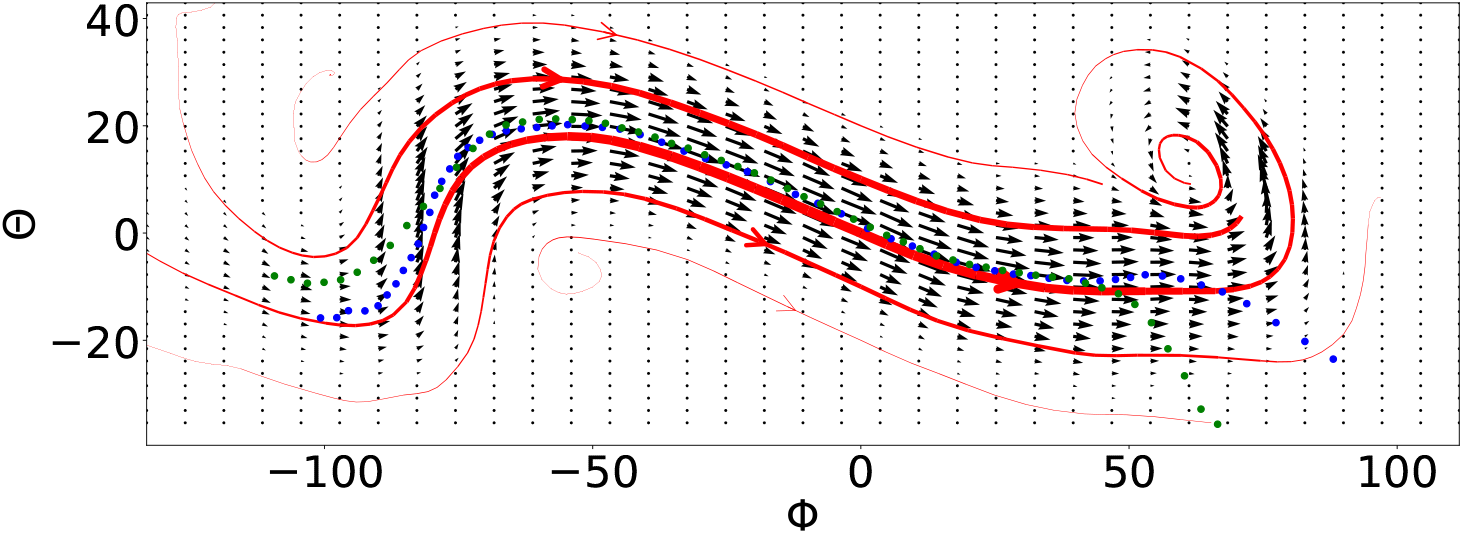
The current, flow lines and principal curves in the plane of two dihedrals *ϕ* and *θ* calculated over the sub-ensemble *PB*-10. The five flow lines pass the points [0,-20], [0,-10], [0,0], [0,10], and [0,20] on the surface *ϕ* = 0.

One may notice that the current vectors near the transition state region or the barrier top in the plane of *ϕ* and *θ* are not parallel but rotated by about 62^*°*^ to the *RC*^*opt*^, a one-dimensional reaction coordinate obtained by maximizing the density-weighted average flux,^63^ which is largely coincide with the one orthogonal to the stochastic separatrix. What makes the maximization of the density-weighted average flux to produce a coordinate whose direction is different to the direction of the reactive current? Since the time-lagged reactive flux for a particular lag time *τ* ∗ may resemble *G*^*U*^ (*ξ′, T*), a quantity closely related to the density-weighted average flux when assuming a uniform average velocity of *ξ*, we thus checked the time-lagged reactive flux along *RC*^*opt*^ for lag times *τ* ranging from 0.5 fs to 1000 fs (Fig. 6A). When *τ* = 200 *fs*, the ratio of average velocity 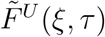 along *RC*^*opt*^ in the range of *RC*^*opt*^ ∈ [−40, 0] remains almost unchanged and is close to 1, while it increases to its maximum at *RC*^*opt*^ = 7^*°*^ and then decreases linearly in the range of *RC*^*opt*^ ∈ [0, 40]. As we know that the barrier top locates at *RC*^*opt*^ = 0, the flat segment of 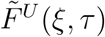 thus corresponds to the stage of climbing up the barrier and the linear decreasing segment corresponds to the one of moving downhill. When moving downhill, a significant portion of the population should be able to commit to the state B within a lag time of 200 fs. That is to say, *τ* = 200 *fs* for the states moving downhill is actually an intermediate time and 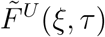 should follow Eq. (29). Thus, the behaviour of the states in the range of *RC*^*opt*^ ∈ [0, 40] is a mixture of a uniform diffusion and the commitment to the state B, and 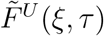 can only be described by a mixture of 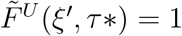 and Eq. (29). The closer the states are to the state B, the higher the portion of the population that commits to the state B is. The states near the barrier top should be largely follow a uniform diffusion. Interesting, the time-lagged fluxes near the barrier top reach their maximum at about *τ* = 200 *fs* (Fig. 6B). The transmission coefficient analogue equals 1 almost everywhere when *τ* = 200 *fs* as well, while it presents a local minimum near the barrier top when *τ* = 0.5 *fs* and it is about 0.43 at *RC*^*opt*^ = 0 (Fig. S3). We thus checked the time-lagged reactive current at *τ* = 200 *fs* in the plane of *ϕ* and *θ* (Fig. 7) and observed that the directions of the time-lagged reactive current near the barrier top follow a vector 0.97***ψ*** + 0.22***θ*** (*α* = 13^*°*^), which is different to the one when *τ* = 0.5 *fs*. Thus, the direction of the time-lagged current is rotating towards the direction of the *RC*^*opt*^ as *τ* increases.

**Figure 6:**
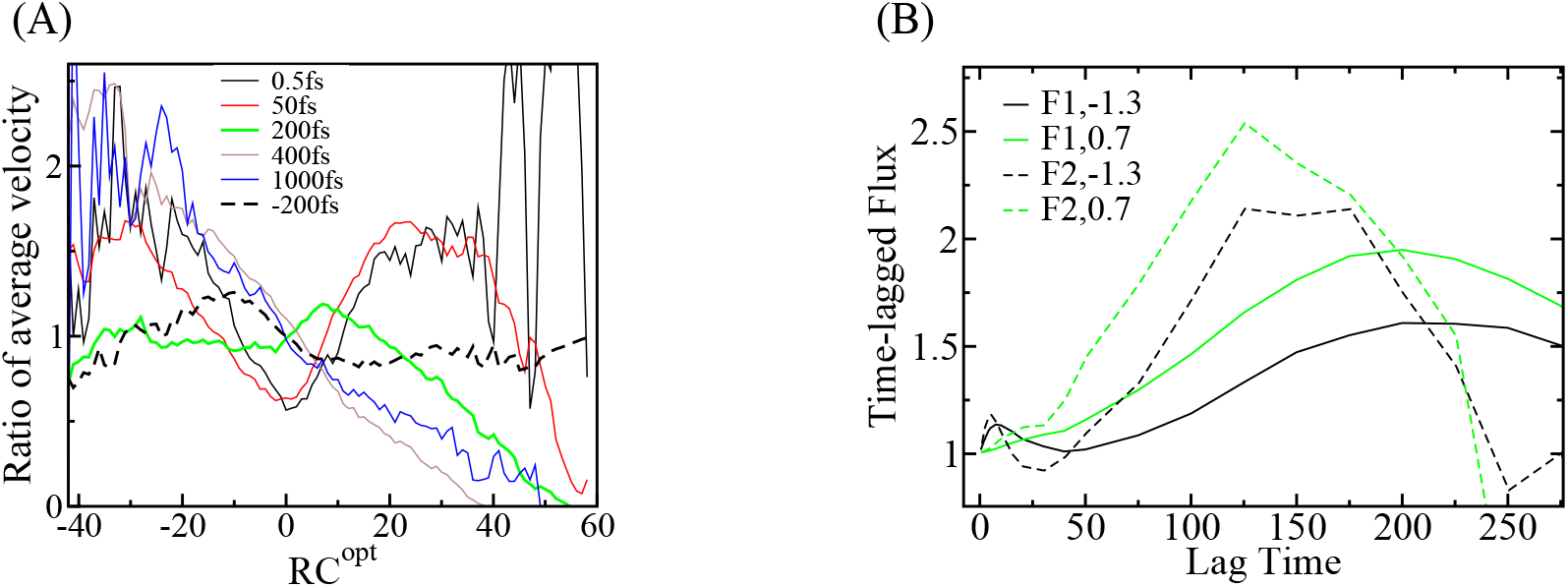
(A) The ratio of average velocity along *RC*^*opt*^ at various lag times. (B) The time-lagged forward flux (F1) and the time-lagged instantaneous flux (F2) (in unit of *N*_*T*_) at points (*RC*^*opt*^ = 1.3^*°*^ and 0.7^*°*^) near the barrier top as a function of the lag time *τ* (in unit of *fs*). The time-lagged forward flux equals the time-lagged instantaneous flux when the time-lagged forward flux reaches its extreme.

**Figure 7:**
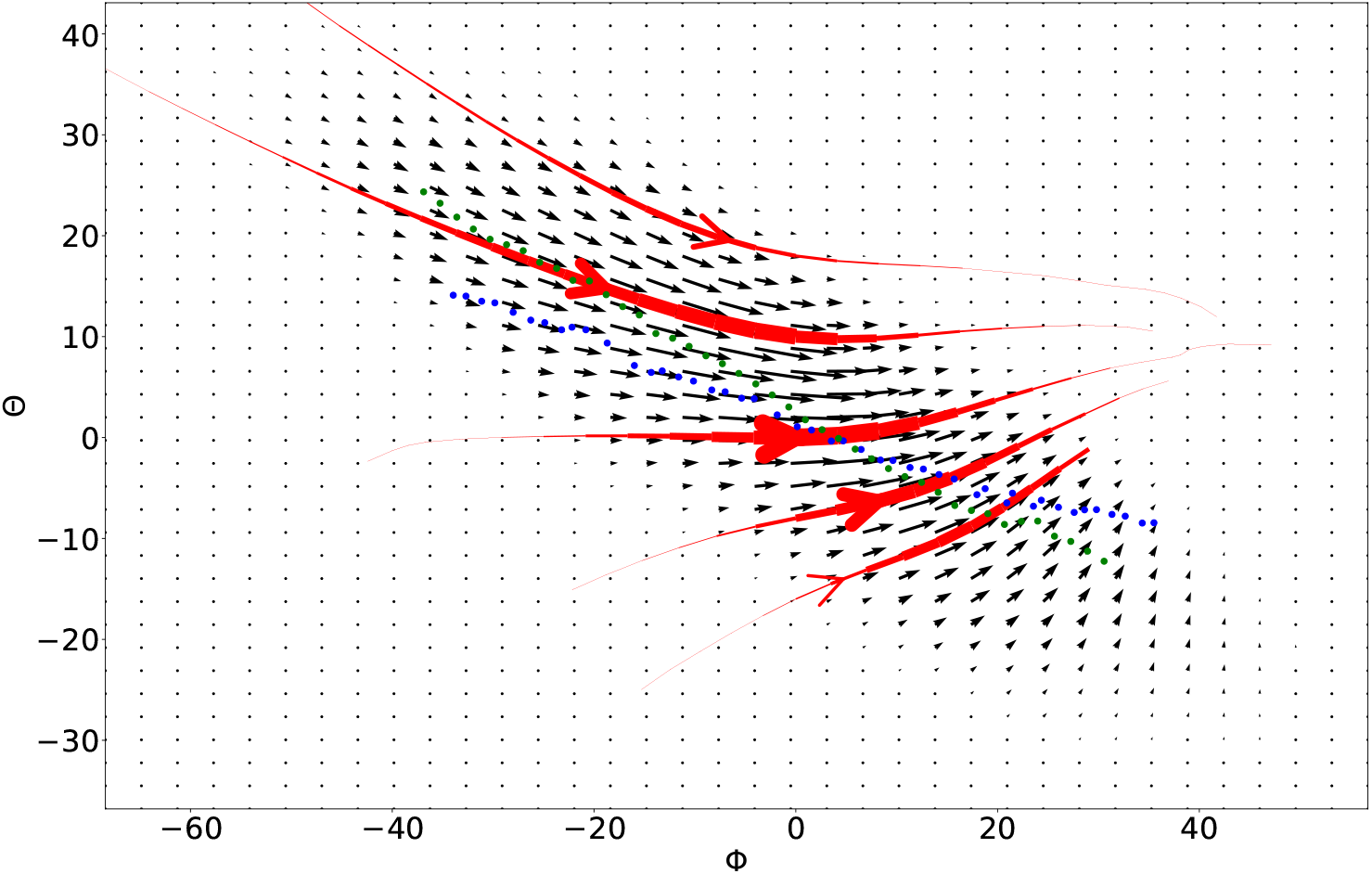
The time-lagged current, flow lines and principal curves for a lag time of *τ* = 200 *fs* in the plane of two dihedrals *ϕ* and *θ* calculated over the sub-ensemble *PB*0. The flow lines pass the same points as the ones in Fig. 2.

Due to the distinct behaviour of 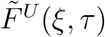 along *RC*^*opt*^ in climbing up the barrier to the one of moving downhill, we thus checked the time-lagged backward reactive flux from B to A for different lag times. In contrast to the case of *τ* = 200 *fs*, we found that 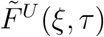 along *RC*^*opt*^ for the backward flux presents a flat region in the range of *RC*^*opt*^ ∈ [5, 50] and increases linearly to its maximum around *RC*^*opt*^ = −10^*°*^ before decreasing to a value around 1 in the range of *RC*^*opt*^ ∈ [−40, 5] when *τ* = −200 *fs* (dashed black line in Fig. 6A). The average direction of the time-lagged reactive current near the barrier top (Fig. S4) is found to be consistent with the one when *τ* = 200 *fs*.

The dependence of the current to the lag time in the plane of *ϕ* and *θ* is generally observed. Interestingly, the forward current is observed to be quite different to the backward current for a lag time of e.g., *τ* = 10 *fs* (Fig. S5), indicating that the projected dynamics in the subspace of *ϕ* and *θ* is not diffusive dynamics and the committor is likely to be dependent on the velocity of these CVs. When the dynamics is projected to the plane of *ϕ* and *ψ*, we did not observe obvious dependence of the current to the lag time and the fluxes at various lag time (up to *τ* = 400 *fs*) in the center of the transition tube largely follow the same path and is very close to the principal curve in the plane.

In Fig. 8, we compared the principal curves obtained in various cases that vary in the subensemble, the one-dimensional reaction coordinate (the coordinate *s*(***y***)), and the lag time. These principal curves are largely consistent. Interestingly, the principal curves obtained along the *RC*^*opt*^ from different sub-ensembles are very close to each other. The principal curve obtained from time-lagged currents for a lag time of *τ* = 200 *fs* largely coincide with other principal curves as well. Thus, a reasonable principal curve can be obtained even when a one-dimensional representation of the reaction coordinate is of moderate quality.

**Figure 8:**
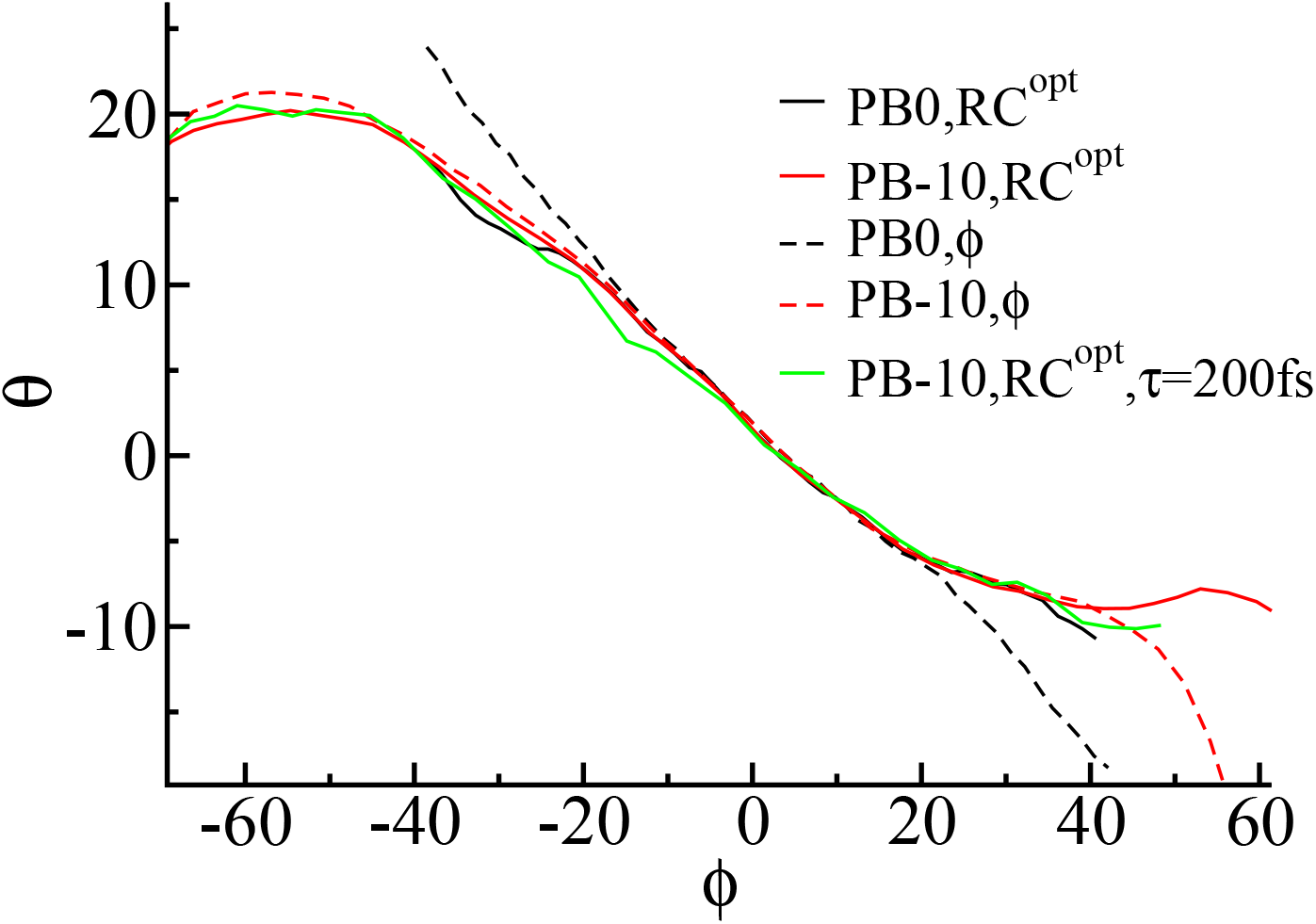
The principal curves in the plane of two dihedrals *ϕ* and *θ* obtained from various cases. Solid (dashed) lines are principal curves obtained with a coordinate *RC*^*opt*^ (*ϕ*) as the coordinate *s*(***y***). Linear regression to the region of *ϕ* ∈ [−15, 10] gives a line 0.49*ϕ* + *θ* = 2.8.

## Concluding Remarks

In order to apply the transition path theory to study complex biomolecular systems, we here generalized the transition path theory to the space of CVs and proposed formulas for quantities such as reactive current and principal curve that can be applied to ensembles of trajectories such as the TPE. The application of these quantities to study the alanine dipeptide in vacuum revealed that the direction of current vectors in the plane of *ϕ* and *θ* is indeed different to the direction of the gradient of the committor, as implied previously for diffusive dynamics.^7,8^ One may then ask why the maximization of a density-weighted average flux produces a coordinate that is largely parallel to the gradient of the committor and does not follow the direction of current vectors. Note that the maximization of the density-weighted average flux is not the same as the maximization of the flux through a dividing surface near the transition state region. Since the fluxes across the dividing surface of both *ϕ* and *RC*^*opt*^ near the barrier top equal the number of transition paths in the TPE, instead of a unique coordinate, the latter way of maximization in the TPE will results in a spectrum of coordinates including the direction of reactive current vectors. From a theoretical viewpoint, we found that the density-weighted average flux resembles time-correlation functions (it is obviously not the classical flux-flux correlation function) and is closely related to the time-lagged flux for a particular lag time, at which the time-lagged average velocity over isosurfaces of the reaction coordinate is invariant. Interestingly, the average direction of time-lagged current vectors in the barrier top follows the direction of the gradient of the committor to a great extent, which provides a satisfying explanation to the above question. We thus believe that the density-weighted average flux is a promising quantity to appraise the relevance of a coordinate to the reaction coordinate and it should be valuable for identifying a few promising CVs to capture the effective dynamics of complex biomolecular systems when MD trajectories of these systems are available.

In the application to the alanine dipeptide in vacuum, the calculated time-lagged reactive current in planes of different pairs of coordinates provided new insights. First of all, when a proper lag time is identified, the corresponding time-lagged reactive current can provide a coordinate that is close to the true reaction coordinate and such results are largely independent to the sub-ensembles. In contrast, the density-weighted average flux requires a sub-ensemble around the transition state region to identify the correct one-dimensional reaction coordinate. On the other hand, such analyses unveiled the characteristics of the dynamics along important coordinates. Here, the reactive current is localized in a narrower transition tube in the plane of *ϕ* and *θ* compared to the one in the plane of *ϕ* and *ψ*, which may be a plausible explanation to the observation that *θ* is more relevant to the reaction coordinate than *ψ*.^46,61–65^ In addition, the direction of the time-lagged reactive current changes significantly and the forward current is not coincident with the backward one in the plane of *ϕ* and *θ*, a phenomenon that is missing in the plane of *ϕ* and *ψ*. Due to the unique behaviour of *θ*, the dynamics projected onto the plane of *ϕ* and *θ* is largely Langevin dynamics and a proper lag time should be chosen to ensure the Markovity of the effective dynamical propagator.

Since the principal curves obtained in various conditions are largely consistent, we expect that a reasonable principal curve can be rather easily obtained once proper CVs are chosen by analysing the density-weighted average flux and the time-lagged reactive current over the TPE. Note that the principal curve is the main target of the string method with swarms-of-trajectories.^8,14^ If such a principal curve is provided, it will be useful for unveiling the detailed reaction mechanism in multiple aspects. For example, it can provide the main shape of the transition tube and the free energy along it can be estimated with extra and affordable simulations. ^13,14,70^

Given that the TPE is generally available for complex biomolecular systems, ^20^ we thus believe that the proposed theory is applicable to these systems. Although we illustrated the application of the proposed theory to the TPE of a simple system, the proposed quantities can be calculated for ensembles other than the TPE, for example an ensemble of non-equilibrium short trajectories that can be analysed by non-equilibrium non-parametric analysis^57^ and Markov state models.^58,59^

## Appendix

### (A) Proof of *F*^*U*^ (*f* (*ξ′*)) = *F*^*U*^ (*ξ′*) and 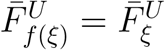

If *f* (*x*) is a monotonically increasing function of *x* and is continuously differentiable, then its derivative *f′* (*x*) *>* 0 everywhere and *x*_0_ is the only root of the function *f* (*x*) − *f* (*x*_0_). From the properties of the Dirac delta distribution, one has

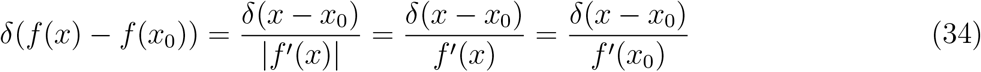

The flux at a give location *f* (*ξ′*) along a generalized coordinate *f* (*ξ*) in an arbitrary ensemble *U* is

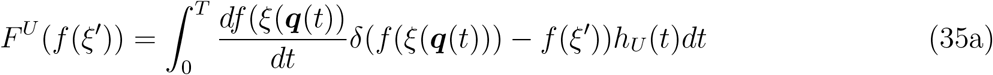

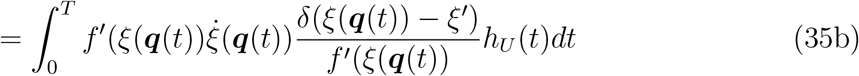

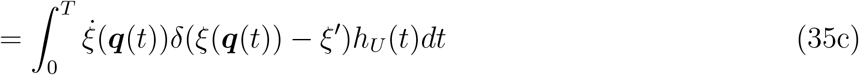

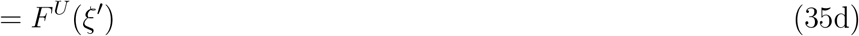

From Eq. (35a) to Eq. (35b), Eq. (34) is used.

The density-weighted average flux of the generalized coordinate *f* (*ξ*) is

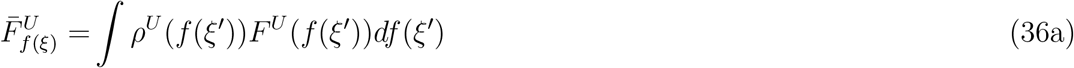

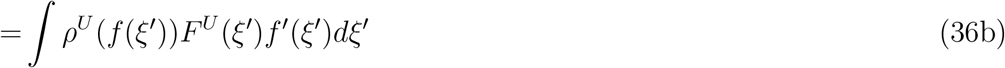

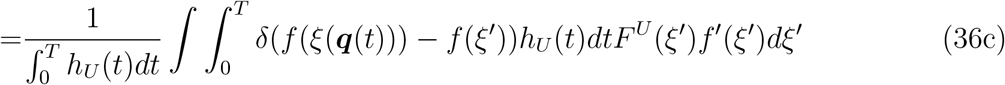

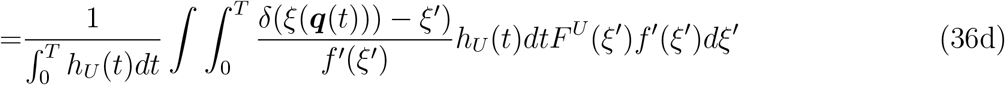

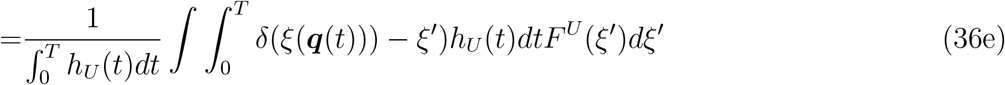

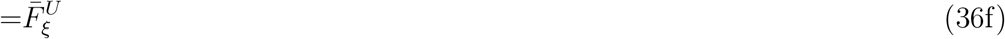

From Eq. (36b) to Eq. (36c), Eq. (10b) is used. From Eq. (36c) to Eq. (36d), Eq. (34) is used.

### (B) The derivation of Eq. (12)

The current in the CVs space is the projection onto it of a non-normalized current in the configuration space 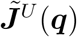, which differs from the probability current in Eq. (3) by a factor of 1*/T* and is given by

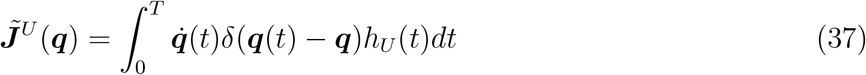

Then, 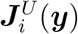, the *i*-th element of ***J***^*U*^ (***y***), is given by

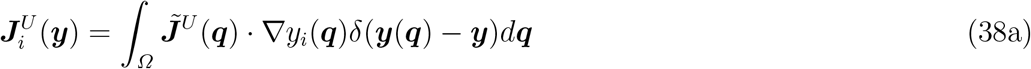

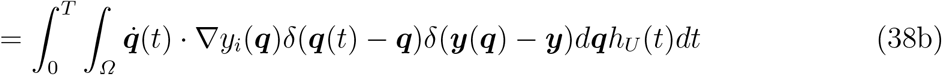

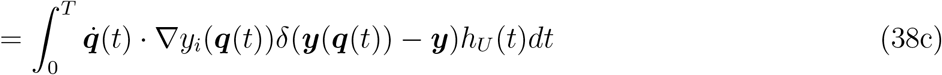

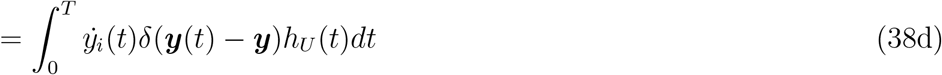

Thus, Eq. (12) is obtained.

## Supplementary Material

See supplementary material for the supplemental Figures S1-S5.

## Acknowledgments

This work was supported by Natural Science Foundation of Guangdong Province, China (Grant No. 2020A1515010984) and the Start-up Grant for Young Scientists (860-000002110384), Shenzhen University.

## References

[1] Chandler, D. The Journal of Chemical Physics 1978, 68, 2959–2970.

[2] Weinan, E.; Vanden-Eijnden, E. Annual review of physical chemistry 2010, 61, 391–420.

[3] Bolhuis, P. G.; Chandler, D.; Dellago, C.; Geissler, P. L. Annual Review of Physical Chemistry 2002, 53, 291–318.

[4] Onsager, L. Physical Review 1938, 54, 554.

[5] Ryter, D. Physica A: Statistical Mechanics and its Applications 1987, 142, 103–121.

[6] Du, R.; Pande, V. S.; Grosberg, A. Y.; Tanaka, T.; Shakhnovich, E. S. The Journal of Chemical Physics 1998, 108, 334.

[7] Weinan, E.; Vanden-Eijnden, E. Journal of statistical physics 2006, 123, 503–523.

[8] Johnson, M. E.; Hummer, G. The Journal of Physical Chemistry B 2012, 116, 8573–8583.

[9] Metzner, P.; Schütte, C.; Vanden-Eijnden, E. Multiscale Modeling & Simulation 2009, 7, 1192– 1219.

[10] Weinan, E.; Ren, W.; Vanden-Eijnden, E. Chemical Physics Letters 2005, 413, 242–247.

[11] Weinan, E.; Ren, W.; Vanden-Eijnden, E. Journal of Chemical Physics 2007, 126, 164103.

[12] Weinan, E.; Ren, W.; Vanden-Eijnden, E. J. Phys. Chem. B 2005, 109, 6688–6693.

[13] Maragliano, L.; Fischer, A.; Vanden-Eijnden, E.; Ciccotti, G. The Journal of chemical physics 2006, 125, 024106.

[14] Pan, A. C.; Sezer, D.; Roux, B. The journal of physical chemistry B 2008, 112, 3432–3440.

[15] Roux, B. The Journal of Physical Chemistry A 2021, 125, 7558–7571.

[16] Bartolucci, G.; Orioli, S.; Faccioli, P. The Journal of chemical physics 2018, 149, 072336.

[17] Metzner, P.; Schütte, C.; Vanden-Eijnden, E. The Journal of chemical physics 2006, 125, 084110.

[18] Karplus, M.; McCammon, J. A. Nature structural biology 2002, 9, 646–652.

[19] Wang, X.; Singh, N.; Li, W. Systems Medicine; Elsevier, 2021; pp 182–189.

[20] Chong, L. T.; Saglam, A. S.; Zuckerman, D. M. Current opinion in structural biology 2017, 43, 88–94.

[21] Jonsson, H.; Mills, G.; Jacobsen, K. W. Classical and Quantum Dynamics in Condensed Phase Simulations); World Scientific, 1998; pp 385–404.

[22] Henkelman, G.; Uberuaga, B. P.; Jónsson, H. The Journal of Chemical Physics 2000, 113, 9901.

[23] Allen, R. J.; Valeriani, C.; ten Wolde, P. R. Journal of Physics: Condensed Matter 2009, 21, 463102.

[24] Borrero, E. E.; Escobedo, F. A. The Journal of chemical physics 2007, 127, 164101.

[25] Faradjian, A. K.; Elber, R. The Journal of chemical physics 2004, 120, 10880–10889.

[26] Elber, R. Quarterly reviews of biophysics 2017, 50.

[27] Berezhkovskii, A. M.; Szabo, A. The Journal of chemical physics 2019, 150, 054106.

[28] Zuckerman, D. M.; Chong, L. T. Annual review of biophysics 2017, 46, 43–57.

[29] Dellago, C.; Bolhuis, P. G.; Csajka, F. S.; Chandler, D. The Journal of Chemical Physics 1998, 108, 1964.

[30] Bolhuis, P. G.; Dellago, C.; Chandler, D. Faraday Discussions 1998, 110, 421–436.

[31] Dellago, C.; Bolhuis, P. G.; Chandler, D. The Journal of Chemical Physics 1999, 110, 6617.

[32] Dellago, C.; Bolhuis, P. G.; Geissler, P. L. Adv. in Chem. Phys. 2002, 123, 1–78.

[33] Li, W.; Gräter, F. Journal of the American Chemical Society 2010, 132, 16790–16795.

[34] Quaytman, S. L.; Schwartz, S. D. Proceedings of the National Academy of Sciences 2007, 104, 12253–12258.

[35] Basner, J. E.; Schwartz, S. D. Journal of the American Chemical Society 2005, 127, 13822– 13831.

[36] Juraszek, J.; Bolhuis, P. G. Proceedings of the National Academy of Sciences 2006, 103, 15859– 15864.

[37] Vreede, J.; Juraszek, J.; Bolhuis, P. G. Proceedings of the National Academy of Sciences 2010, 107, 2397–2402.

[38] Best, R. B.; Hummer, G. Proceedings of the National Academy of Sciences of the United States of America 2005, 102, 6732–6737.

[39] Hu, J.; Ma, A.; Dinner, A. R. Proceedings of the National Academy of Sciences 2008, 105, 4615–4620.

[40] Knott, B. C.; Haddad Momeni, M.; Crowley, M. F.; Mackenzie, L. F.; Götz, A. W.; Sandgren, M.; Withers, S. G.; Ståhlberg, J.; Beckham, G. T. Journal of the American Chemical Society 2013, 136, 321–329.

[41] Bolhuis, P. G. Proceedings of the National Academy of Sciences 2003, 100, 12129–12134.

[42] Best, R. B.; Hummer, G. Proceedings of the National Academy of Sciences 2016, 113, 3263– 3268.

[43] Li, W.; Ma, A. Molecular simulation 2014, 40, 784–793.

[44] Berezhkovskii, A.; Szabo, A. The Journal of chemical physics 2005, 122, 014503.

[45] Rhee, Y. M.; Pande, V. S. The Journal of Physical Chemistry B 2005, 109, 6780–6786.

[46] Ma, A.; Dinner, A. R. J Phys Chem B 2005, 109, 6769–79.

[47] Peters, B.; Trout, B. L. The Journal of Chemical Physics 2006, 125, 054108.

[48] Peters, B.; Beckham, G. T.; Trout, B. L. The Journal of chemical physics 2007, 127, 034109– 034109.

[49] Peters, B.; Bolhuis, P. G.; Mullen, R. G.; Shea, J.-E. The Journal of chemical physics 2013, 138, 054106.

[50] Mori, Y.; Okazaki, K.-i.; Mori, T.; Kim, K.; Matubayasi, N. The Journal of Chemical Physics 2020, 153, 054115.

[51] Antoniou, D.; Schwartz, S. D. The Journal of chemical physics 2009, 130, 151103.

[52] Antoniou, D.; Schwartz, S. D. The Journal of Physical Chemistry B 2011, 115, 2465–2469.

[53] Brandt, S.; Sittel, F.; Ernst, M.; Stock, G. The journal of physical chemistry letters 2018, 9, 2144–2150.

[54] Jung, H.; Covino, R.; Hummer, G. arXiv preprint 1901.04595 2019,

[55] Li, W.; Ma, A. The Journal of chemical physics 2015, 143, 11B603 1.

[56] Elber, R.; Bello-Rivas, J. M.; Ma, P.; Cardenas, A. E.; Fathizadeh, A. Entropy 2017, 19, 219.

[57] Krivov, S. V. J. Chem. Theory Comput. 2021, 17, 5466–5481.

[58] Chodera, J. D.; Nóe, F. Current opinion in structural biology 2014, 25, 135–144.

[59] Wu, H.; Nüske, F.; Paul, F.; Klus, S.; Koltai, P.; Nóe, F. The Journal of chemical physics 2017, 146, 154104.

[60] Hummer, G. The Journal of chemical physics 2004, 120, 516.

[61] Li, W.; Ma, A. The Journal of chemical physics 2016, 144, 134104.

[62] Li, W. The Journal of chemical physics 2018, 148, 084105.

[63] Li, W. The Journal of Chemical Physics 2022, 156, 054117.

[64] Bolhuis, P. G.; Dellago, C.; Chandler, D. Proceedings of the National Academy of Sciences 2000, 97, 5877–5882.

[65] Li, W.; Ma, A. The Journal of Chemical Physics 2016, 144, 114103.

[66] Tromp, J. W.; Miller, W. H. Faraday Discussions of the Chemical Society 1987, 84, 441–453.

[67] Thompson, W. H.; Miller, W. H. The Journal of chemical physics 1997, 106, 142–150.

[68] Goussev, A.; Schubert, R.; Waalkens, H.; Wiggins, S. The Journal of chemical physics 2010, 133, 244113.

[69] Tromp, J. W.; Dumont, R. S. Molecular Physics 2012, 110, 817–824.

[70] Vanden-Eijnden, E.; Venturoli, M. The Journal of Chemical Physics 2009, 130, 194103.

