## supplemental Figures S1-S5 for "Time-lagged Flux in the Transition Path Ensemble: Flux Maximization and Relation to Transition Path Theory"

Wenjin Li
Institute for Advanced Study, Shenzhen University, Shenzhen, China

### Supplementary Material

#### Contents:

- Supplemental Figures S1-S5.

### Supplemental Figures

S1 Probability Densities of Sub-ensembles . . . . . . . . . . S2
S2 Flux Along $\phi$ . . . . . . . . . . S2
S3 Transmission Coefficient Analogue . . . . . . . . . . S3
S4 The Time-lagged Backward Current at $\tau = 200\,fs$ . . . . . . . . . . S4
S5 The Time-lagged Forward and Backward Current at $\tau = 10\,fs$ . . . . . . . . . . S5

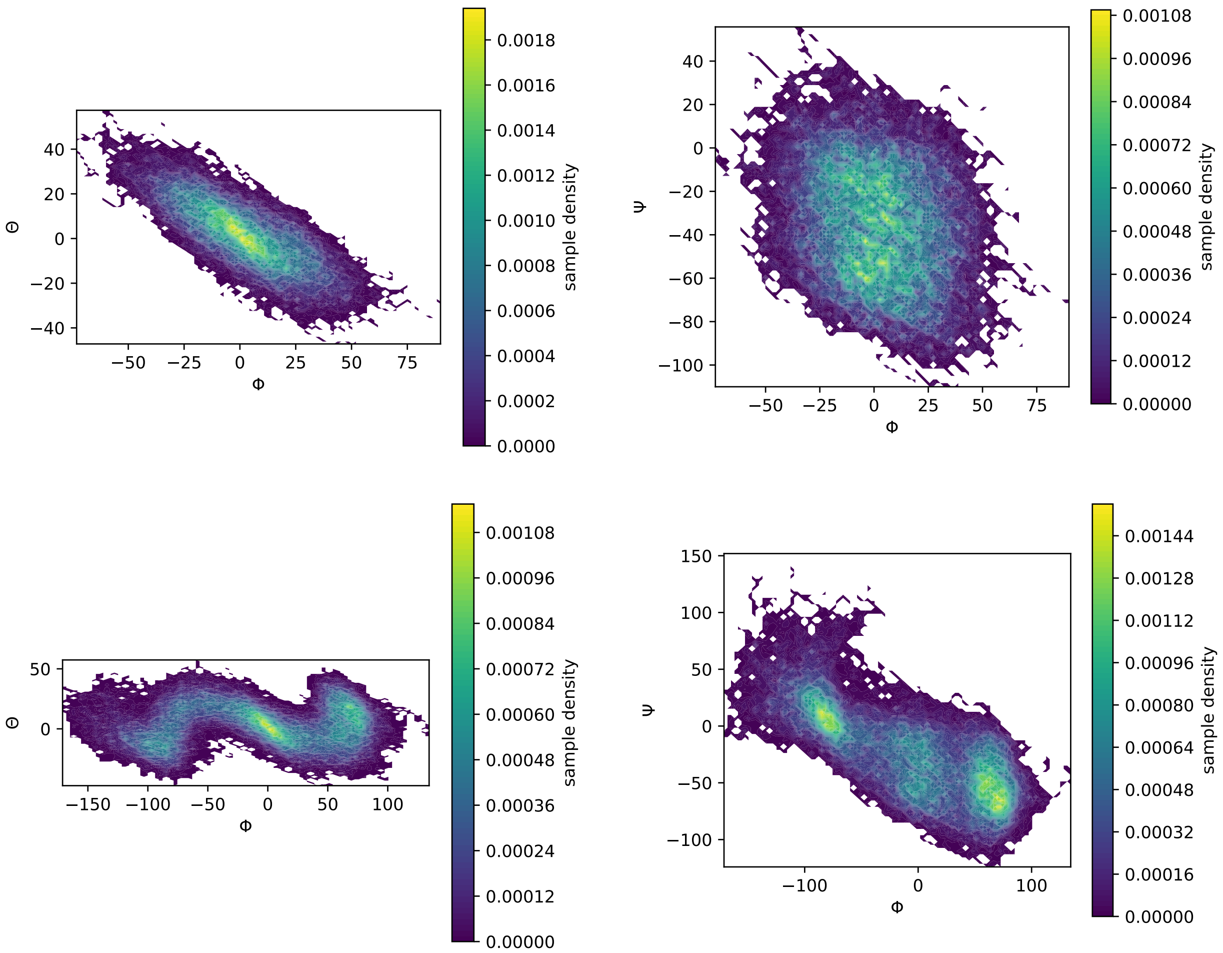


Figure S1: The probability densities of A-B reactive trajectories projected on the plane of $\phi$ and $\theta$ (Left) or the plane of $\phi$ and $\psi$ (Right) in the sub-ensemble $PB0$ (Top) and the sub-ensemble $PB$-10 (Bottom).

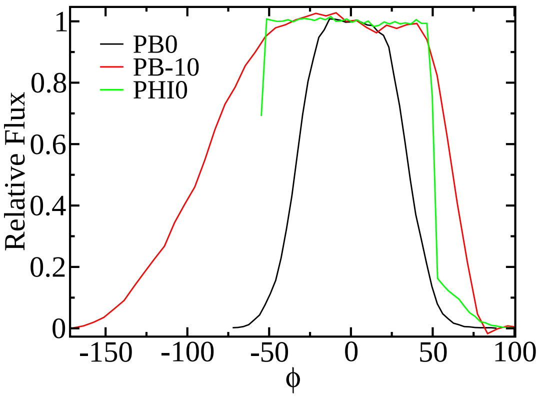


Figure S2: The relative flux along $\phi$ at different sub-ensembles. The relative flux is the flux divided by the number of transition paths in the transition path ensemble.

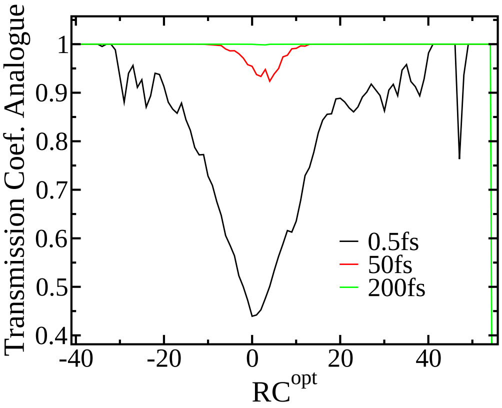


Figure S3: The time-lagged transmission coefficient analogues along the $RC^{opt} = 0.83\phi + 0.56\theta$ at different lag times.

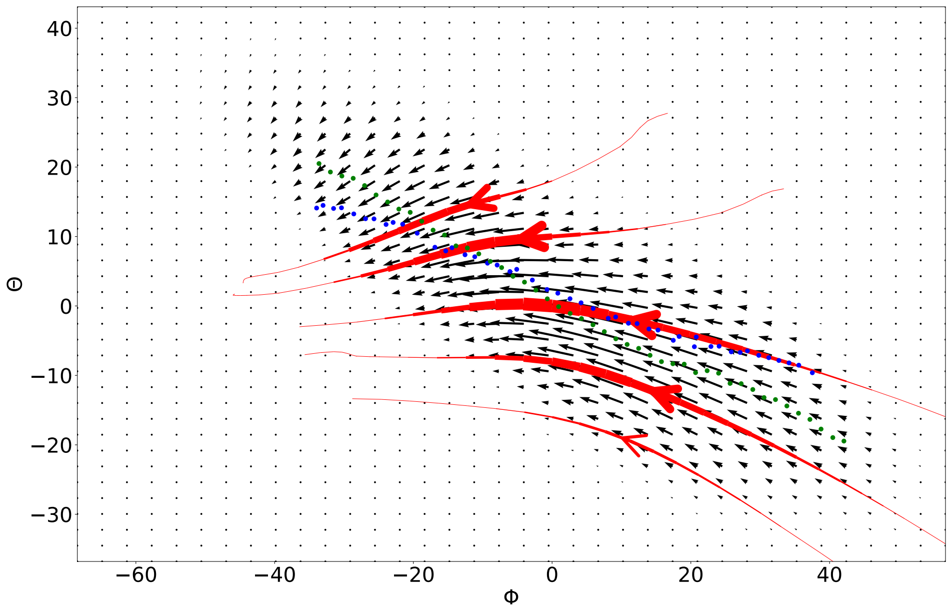


Figure S4: The time-lagged backward current, flow lines and principal curves for a lag time of $\tau = 200\,fs$ in the plane of two dihedrals $\phi$ and $\theta$ calculated over the sub-ensemble $PB0$. Five flow lines in red pass the points [0,-16], [0,-8], [0,0], [0,10], and [0,18] on the surface $\phi = 0$. The width of a flow line varies with the magnitude of the current vector. Blue and green dotted lines are two principal curves obtained with a coordinate $RC^{opt} = 0.83\phi + 0.56\theta$ and $\phi$ as the coordinate $s(\boldsymbol{y})$, respectively. The units of both axes are degree.

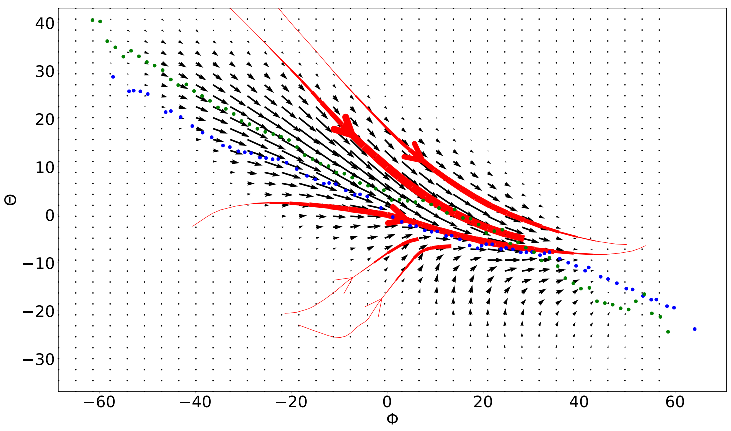


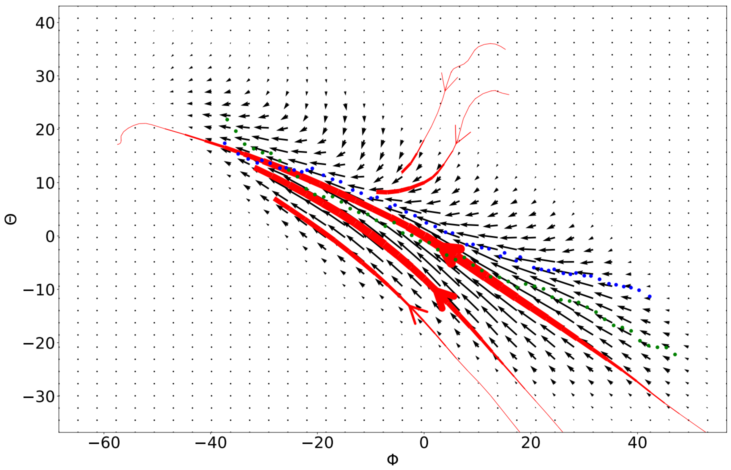


Figure S5: The time-lagged forward (Top) and backward (Bottom) current, flow lines and principal curves for a lag time of $\tau = 10\,fs$ in the plane of two dihedrals $\phi$ and $\theta$ calculated over the sub-ensemble $PB0$.
